# Interlocked transcription factor feedback loops maintain and restore touch sensation

**DOI:** 10.1101/2025.05.15.654349

**Authors:** Filipe Marques, Yihan Chen, Honorine Destain, Margaux Marinelli, Paschalis Kratsios

## Abstract

The sense of touch relies on the continuous function of specialized mechanosensory circuits, but the underlying molecular mechanisms remain poorly understood. Here, we report that the conserved transcription factors (TFs) CFI-1 (ARID3) and EGL-5 (HOXA7) jointly maintain in adult *C. elegans* the molecular identity of two key interneuron types, securing information processing within a mechanosensory circuit. Toggling between normal and low levels of CFI-1 or EGL-5 in adults generated digital-like (ON/OFF) effects both on touch-evoked escape response and interneuron identity. Strikingly, reintroduction of CFI-1 following its prolonged depletion restored escape response defects. Mechanistically, we identified two network motifs, a double-positive CFI-1/EGL-5 feedback loop and positive CFI-1 autoregulation, which together “lock-in” the interneuron identity programs. We propose that these interlocked motifs not only maintain robust escape responses throughout life, but are also essential for the restorability of adult-onset touch defects. Altogether, this work illuminates the molecular principles that maintain adult neuron identity and circuit function, and offers biomedically relevant insights into the restorability of neuronal and behavioral defects caused by mutations or variation in TF-encoding genes.

## INTRODUCTION

Mechanosensation is an ancient sensory modality that allows animals to perceive and interact with their environment^1^. By transducing mechanical stimuli into neuronal signals, it provides the basis for the senses of touch, hearing, proprioception, and pain. Importantly, various human conditions impair mechanosensation ^2^ ^3^ ^4^. Yet, our understanding of the molecular mechanisms that establish functional mechanosensory circuits during development or maintain them throughout adulthood remains limited, especially compared to other well-studied sensory (e.g., visual) circuits ^5^ ^6^. This knowledge gap is partly due to the inherent complexity of vertebrate nervous systems ^7^ ^6^ ^8^. Studying how mechanosensory circuits become and remain functional can help us understand how organisms sustain the ability to properly respond to mechanical stimuli, such as touch-evoked escape responses, over their lifetime.

A fundamental organizational principle of mechanosensory circuits is their three-tiered structure: (a) specialized mechanosensory neurons in the peripheral nervous system, known as mechanoreceptors, detect and convert mechanical stimuli into electrical signals, (b) interneurons in the central nervous system (CNS) then integrate and process this information, and finally (c) motor neurons convey commands to peripheral muscles, driving appropriate behavioral responses.

Research on mechanosensory circuits started almost a century ago ^9^, and has primarily focused on mechanosensory and motor neurons ever since, largely due to their experimental accessibility. At the mechanosensory level, significant progress has been made in discovering ion channels that detect mechanical stimuli ^10^ ^11^ ^12^, and in identifying transcriptional mechanisms that govern mechanosensory neuron development across species ^13^ ^14^ ^15^ ^16^ ^17^ ^18^. Notably, members of the POU family of homeodomain transcription factors (TFs) control mechanosensory neuron identity in nematodes (*C. elegans*) ^19^ ^20^ ^21^ ^22^ ^23^ ^24^), sea anemones (*Nematostella vectensis*) ^25^ ^26^, and mice (*M. musculus*) ^27^. Similarly, our understanding of transcriptional programs at the motor level is also extensive; numerous studies have identified conserved roles for HOX and LIM homeodomain TFs in motor neuron development ^28–30^. In contrast, the genetic programs that control the development and function of interneurons remain poorly understood in all model systems ^31^, as these cells are often experimentally inaccessible due their CNS location.

The sustained function of any neuronal circuit relies on the ability of its constituent neuron types to continuously express distinct sets of terminal identity genes, encoding essential neuronal proteins (e.g., neurotransmitter [NT] receptors, NT biosynthesis, gap junction proteins, ion channels, neuropeptides)^32^. Studies across model organisms have identified specific transcription factors (TFs), called “terminal selectors”, that induce during development the expression of terminal differentiation genes in specific neuron types^33–35^. To date, dozens of terminal selectors have been described in multiple animal species, including nematodes, fruit flies, planarians, cnidarians, marine chordates, zebrafish, and mice^36–38^. This widespread presence underscores the deeply conserved role of terminal selectors as critical regulators of neuronal identity and function. Importantly, human genetic studies strongly indicate that numerous terminal selector orthologs play causal roles in neurodevelopmental and neurodegenerative disorders^39–46^. Recent studies indicate that a combination of several TFs can function as terminal selectors in a specific neuron type^47,48^, highlighting the complexity of the underlying transcriptional networks. Yet, the organization and patterns of TF interactions within such networks remain largely unknown.

Terminal selectors are thought to be continuously required for maintenance of neuronal terminal identity throughout life^36^. In mice, inducible genetic approaches have firmly established a maintenance requirement for two terminal selectors, *Nurr1* and *Pet-1/FEV,* in dopaminergic and serotonergic neurons, respectively^49,50^. In *C. elegans,* the advent of auxin-inducible degron ^51^ technology has uncovered maintenance roles for various terminal selectors both on neuron type identity and animal behavior ^52,53^. For example, the POU homeodomain terminal selector UNC-86 is specifically required to maintain mechanosensory neuron identity and normal touch responses ^54^. The terminal selector UNC-3/EBF is required to maintain motor neuron identity and normal locomotion ^55^. Together with other studies^39,56^, these vertebrate and invertebrate examples firmly establish that - in the adult - reduced expression or activity of a terminal selector significantly impairs the terminally differentiated state of a neuron, ultimately impacting organismal behavior. However, whether adult neurons can revert from the impaired (OFF) state back to a fully differentiated (ON) state remains an outstanding question with important clinical implications, as reduced expression or activity of terminal selectors (e.g., *NURR1, FEV*) is linked to neurological conditions in humans, including Parkinson’s disease and autism ^40,57^. In general, our mechanistic understanding of how terminal selectors maintain the identity program of adult neuron types remains poor, preventing our ability to understand and/or restore neuronal defects seen in patients with neurological diseases.

To unveil the mechanisms that maintain adult neuron function and animal behavior, we use as a paradigm a *C. elegans* mechanosensory circuit necessary for touch-evoked escape responses ^12,58,59^. We report that two conserved TFs, CFI-1 (human ARID3) and EGL-5 (human HOXA7), jointly function as terminal selectors in two lumbar interneuron types (PVC and LUA) to secure information processing within this circuit. By integrating extensive genetic analyses with chromatin immunoprecipitation and sequencing (ChIP-Seq) datasets, we propose that CFI-1 and EGL-5 act directly to induce during development, and maintain in adulthood, a broad spectrum of PVC and LUA terminal identity features, including genes involved in NT biosynthesis, gap junction formation, and neuropeptide signaling. Next, we leveraged the AID system, enabling inducible and reversible TF depletion in adult neurons coupled with behavioral analysis. We discovered that adult PVC and LUA interneurons can revert from an impaired OFF state back to the terminally differentiated ON state. Resupply of CFI-1 following its prolonged depletion restored touch defects, a finding with important implications for adult-onset neurological disorders. Mechanistically, we identified two network motifs, a double-positive CFI-1/EGL-5 feedback loop and CFI-1 positive autoregulation, which “lock-in” the interneuron identity programs, and thereby ensure the robustness of touch-evoked escape responses. Altogether, our findings advance our mechanistic understanding of how the terminally differentiated state of a neuron is maintained in adulthood. As CFI-1 and EGL-5 orthologs are expressed in the nervous system of other species^28,60–62^, the molecular principles described herein may be conserved.

## RESULTS

### *cfi-1/ARID3* is selectively required for posterior touch in *C. elegans*

To investigate how TFs control neuronal identity in mechanosensory circuits, we focused on *cfi-1,* the sole *C. elegans* ortholog of the ARID3 family of TFs^63,64^ (**Fig. 1A**). Prior work showed that *cfi-1* is required for normal posterior responses to gentle (light) touch ^65, 66^, but the underlying cellular and molecular mechanisms remained unclear. Because the neuronal circuits that control gentle and harsh touch differ (**Fig. 2C**)^67^, we wondered whether *cfi-1* also controls harsh touch responses. We therefore conducted both gentle and harsh touch assays on wild-type ^68^ and mutant animals carrying either of two *cfi-1* loss-of-function (LOF) alleles, *ky651* or *ot786* (**Fig. 1A**) ^69^ ^65^. Both WT and *cfi-1* LOF mutant animals exhibited normal anterior touch (gentle or harsh) responses (**Fig. S1A-B**). In contrast, responses to posterior touch (both gentle and harsh) were severely impaired in *cfi-1* mutants (**Fig. 1B-C, S1C**). Notably, the *cfi-1* mutant phenotype in both posterior touch assays was not fully penetrant, and can be categorized into three groups: a) animals that responded normally and moved forward (“forward” group), b) animals that did not respond (“no response” group), and c) animals that moved backward (“backward” group) (**Fig. 1B-C**). In these assays, 42.5% of *cfi-1 (ot786)* mutants responded normally (forward group) to gentle touch (**Fig. 1B**), while 23.1% of *cfi-1(ot786)* mutants responded normally to harsh touch (**Fig. 1C**). We conclude that *cfi-1* is selectively required for posterior, but not anterior touch responses. Because the harsh touch defects are stronger (**Fig. 1B-C**), we conducted only harsh touch assays hereafter.

**Figure 1.**
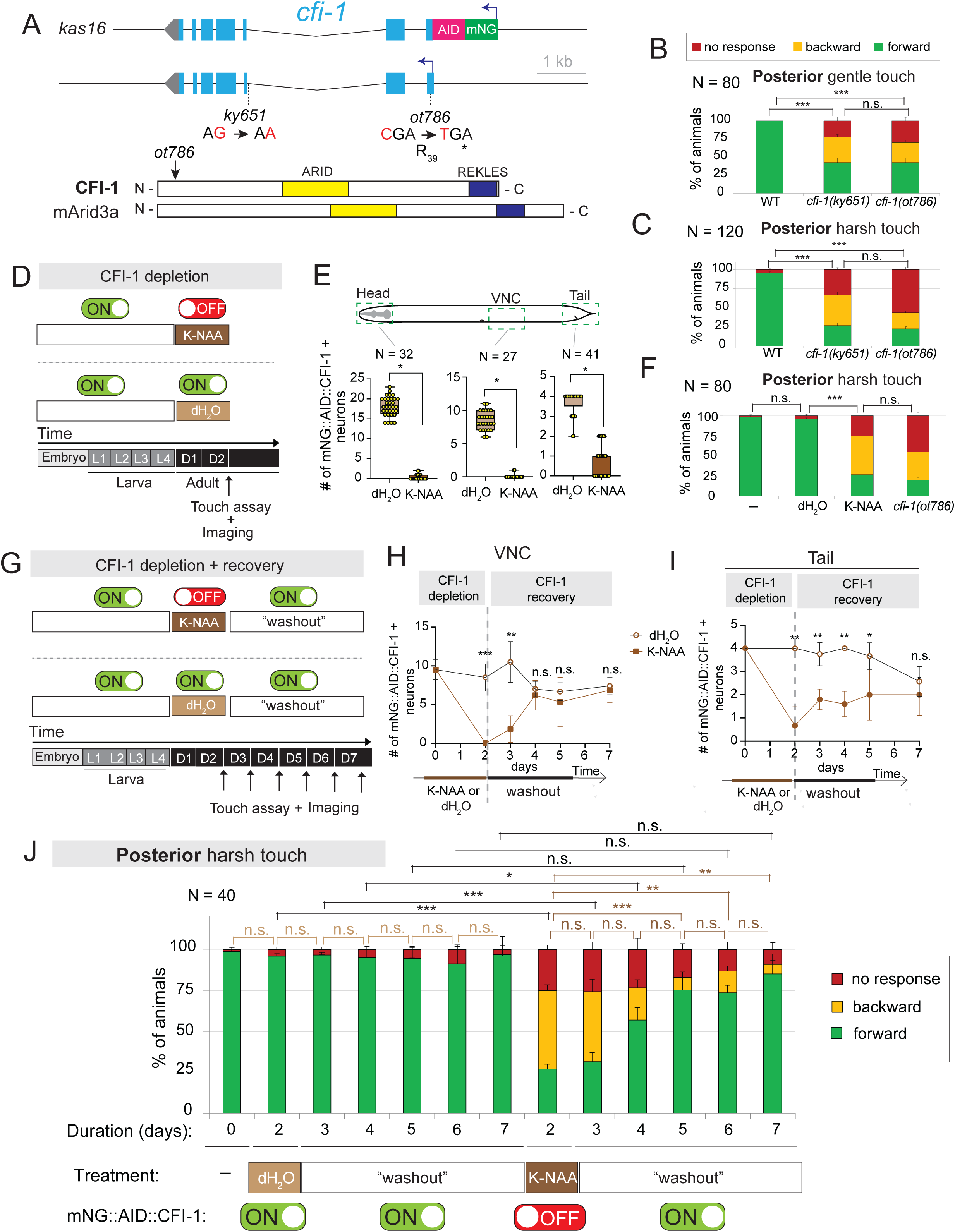
CFI-1 functions as an on-off switch that regulates posterior touch behavior in adult stage. **A.** Diagram of the *cfi-1* locus. *ky651* allele: G-to-A splice acceptor mutation^65^; *ot786* allele: a C-to-T mutation generates a premature STOP^69^; *kas16* allele: mNG::AID::*cfi-1* endogenous reporter; Schematic of CFI-1 protein and its mouse ortholog Arid3a. **B-C.** Posterior gentle (N = 80) and harsh (N = 120) touch assays on day 1 adult control (N2) and *cfi-1* mutant animals. **D**. Diagram of control (dH_2_O) and auxin (K-NAA) treatment for CFI-1 depletion. **E**. Lateral worm view with dashed boxes highlighting the regions where *cfi-mNG::AID::CFI-1*-expressing neurons were counted. Quantifications after the dH_2_O (control) and auxin (K-NAA) treatments in *kas16[mng::aid::cfi-1]; ieSi57 [peft-3::TIR1::mruby::unc-54 3’utr + cbr-unc-119(+)]* animals. N ≥ 27 animals. **F**. Posterior harsh touch assay on day 2 adult animals with no treatment (-), control (dH_2_O) and auxin (K-NAA), as well as *cfi-1(ot786)* mutants for comparison. N = 80 animals. **G**. Diagram of control (dH_2_O) and auxin (K-NAA) treatment for CFI-1 depletion followed by “washout”. **H-I.** Quantification of VNC and tail neurons expressing the endogenous *cfi-1* reporter after water (control), auxin (K-NAA) and “washout” recovery treatments on kas16[mNG::AID::cfi-1]; ieSi57 [eft-3p::TIR1::mRuby::unc-54 3’UTR + Cbr-unc-119(+)] animals. N ≥ 5 animals. **J.** Posterior harsh touch assays with no treatment, and after dH_2_O (control), auxin (K-NAA) and “washout” recovery treatments. N = 40 animals. *Behavioral data in panels B–C, F, and J:* Significance was determined using Bonferroni post-hoc tests. Statistical thresholds are indicated as follows: *p* = 0.0019 (*), p = 0.0009 (**),* p < 0.0001; n.s., not significant. *Panels D, G, and J:* Green (ON) symbols indicate the presence of CFI-1 expression; red (OFF) symbols indicate its absence. *Panel E:* Box-and-whisker plots display the number of neurons expressing the fluorescent reporter. Whiskers represent the minimum and maximum values; the horizontal line within each box denotes the median. Each dot represents the average number of neurons in an individual animal. An unpaired, two-sided *t*-test with Welch’s correction was used: \*\**p* < 0.0001. *Quantifications in panels H–I:* Bonferroni post-hoc tests were used to determine statistical significance: *p* = 0.0082 (*), p = 0.0006 (**),* p = 0.00008.

**Figure 2.**
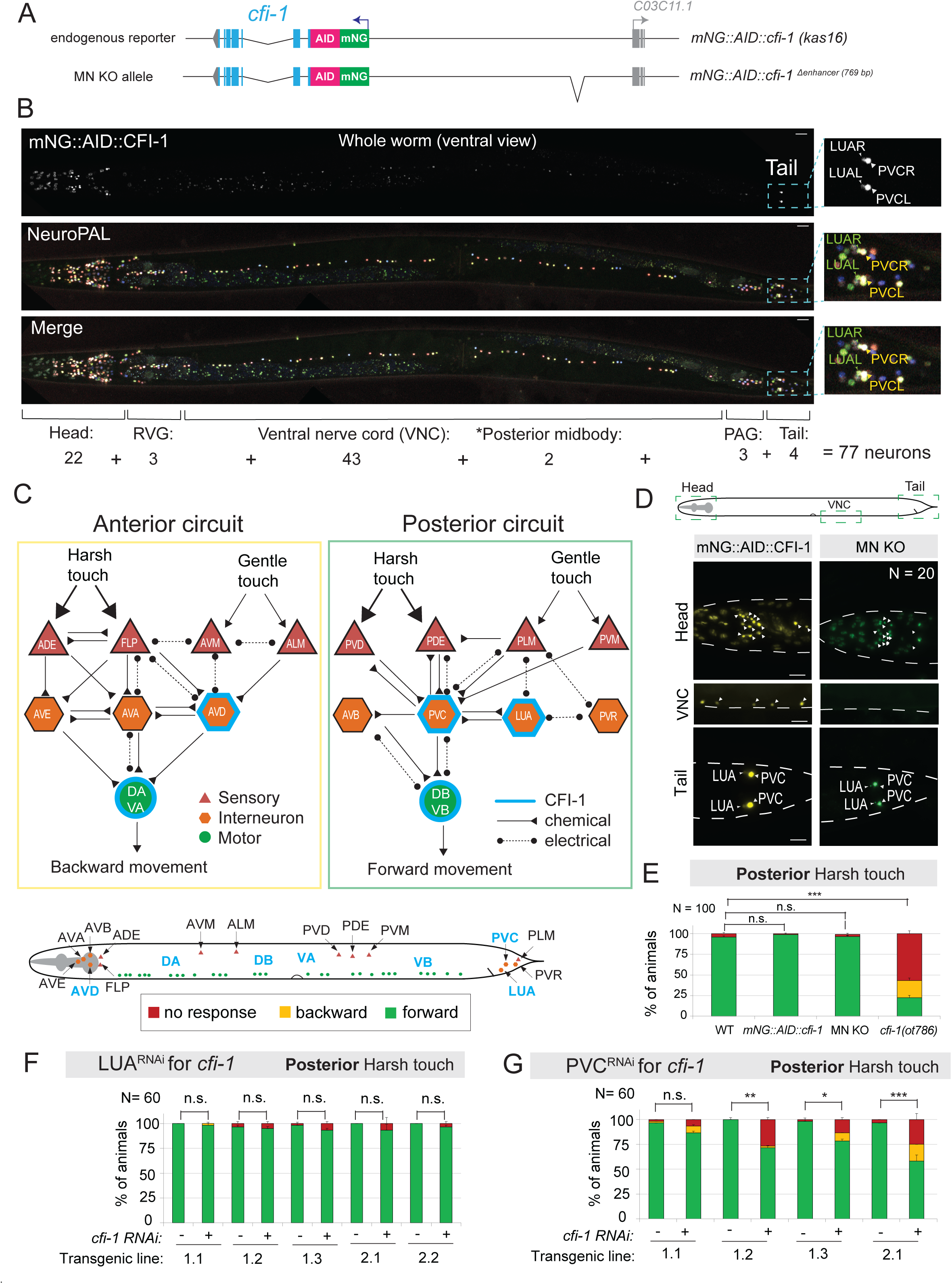
A nervous system-wide CFI-1 expression map and a cell-autonomous CFI-1 role in PVC interneurons. **A.** Diagram of *cfi-1* locus. The MN KO allele carries a 769 bp deletion (-11,329 bp to -12,097 bp) in an enhancer element required for MN expression ^55^. **B.** Representative images of the whole worm (day 1 adult) showing mNG::AID::CFI-1 expression (top) NeuroPAL transgene color code (middle), and merge (bottom) strain. See Table S1. Tail insets are magnified to show PVC and LUA interneurons. N = 3 animals. Scale bars: 20 μm. **C.** Schematic of *C. elegans* mechanosensory circuit ^132^. Circuit assembled using data from nemanode.org. Dataset used: complete adult animal^117^; Connections between neurons have at least 1 chemical synapse or gap junction). For simplicity, not all connections between sensory and/or interneurons are shown. Bottom schematic: cell body position of neurons in the mechanosensory circuits. **D.** Dashed boxes highlight regions shown in images below. Representative images showing the expression of mNG::AID::CFI-1 in head, VNC and tail of *C. elegans* animals carrying the *mNG::AID::cfi-1* and MN KO allele. Arrowheads: neuronal nuclei. N = 20 animals. Scale bars: 20 μm. **E.** Posterior harsh touch assays on animals carrying the mNG::AID::*cfi-1* and MN KO alleles. N = 100 animals. **F.** Posterior harsh touch assay in control and LUA-specific *cfi-1* RNA*i* in young adult animals. Five independent lines tested. N = 60. **G.** Posterior harsh touch assay in *C. elegans* control vs PVC -specific *cfi-1* RNA*i* in young adult animals. Four independent lines were tested. N = 60 animals. For behavior in panels **E-G,** *, p = 0.0093; **, p = 0.0009; ***, p < 0.0001; n.s., not significant, as determined by Bonferroni post-hoc tests.

### CFI-1 functions as an “on-off” switch to maintain and restore posterior touch defects

Because the behavioral assays were conducted on animals constitutively lacking *cfi-1* since early development (**Fig. 1A-C**), it remained unclear whether CFI-1 is continuously required in the adult to maintain normal touch responses. We therefore employed the AID system to induce CFI-1 depletion in adult animals^52^, leveraging an endogenous *cfi-1* reporter allele (*kas16*) tagged with the AID degron (*mNG::AID::cfi-1)*^70^ (**Fig. 1A**). Starting at the end of the fourth larval stage (L4), when the *C. elegans* nervous system is mostly developed, we continuously exposed animals to synthetic auxin (K-NAA) or dH_2_O (control) for two days (**Fig. 1D**). We observed robust depletion of mNG::AID::CFI-1 in adult (day 2) animals in known sites of CFI-1 expression ^65,70^, including head muscle and neurons of the head, ventral nerve cord, and tail (**Fig. 1E, S1G**). Similar to *cfi-1(ot786)* LOF mutants, K-NAA-treated *cfi-1(kas16[mNG::AID::cfi-1])* animals exhibited a significant impairment in posterior harsh touch responses compared to dH_2_O-treated controls (**Fig. 1F**). We obtained similar results when *cfi-1(kas16[mNG::AID::cfi-1])* animals were exposed to natural auxin (IAA) for two days starting at L4 (**Fig. S1H-J**). We conclude that CFI-1 is continuously required in early adulthood for normal touch-evoked escape responses.

Next, we leveraged the reversibility of the AID system to investigate whether the detrimental effects on posterior touch upon adult-specific CFI-1 depletion can be restored when CFI-1 is resupplied. After a two-day exposure to K-NAA or dH_2_O (control), we again observed a robust reduction in the number of mNG::AID::CFI-1-expressing neurons (**Fig. 1H-I, S1G**). Next, we transferred adult (day 3) *cfi-1(kas16[mNG::AID::cfi-1])* animals to regular NGM plates for five days (D3-D7) to “washout” K-NAA (**Fig. 1G, S1E-F**), and observed a gradual and complete recovery by D7 in the number of mNG::AID::CFI-1-expressing neurons (**Fig. 1H-I**, **S1D-G**). Remarkably, the recovery of CFI-1 expression coincided with a progressive restoration of posterior harsh touch defects: 84.8% of animals respond normally at D7 compared to 26.8% at D2 (**Fig. 1J**). Altogether, these findings suggest that CFI-1 operates as a functional “on-off” switch for adult posterior touch responses. When CFI-1 is expressed (ON), the response is normal. When CFI-1 is depleted (OFF), the response is impaired. Flipping the switch from OFF to ON significantly restored the harsh touch defects, indicating that behavioral defects with adult onset can be restored through resupply of a single TF.

### A complete map of CFI-1 expression in *C. elegans*

In which cell types of the *C. elegans* nervous system is *cfi-1* acting to control posterior touch behavior? To address this, we first built a comprehensive expression map using our endogenous *cfi-1(kas16[mNG::AID::cfi-1])* reporter allele crossed to NeuroPAL, a multicolor strain suitable for nervous system-wide cell identification^71^. We found that mNG::AID::CFI-1 is expressed in 77 neurons at the adult (day 1) stage (**Fig. 2B**, **S3I, Table S1**). We obtained similar results with another endogenous reporter allele, *cfi-1* (*dev183[mNG::cfi-1])*^72^ (**Fig. S2B-C**). We further validated the CFI-1 expression pattern by using molecular markers for various neurotransmitter identities (cholinergic, GABAergic, glutamatergic, and dopaminergic) (**Fig. S3**), and analyzing available single-cell RNA sequencing (sc-RNA-Seq) data derived from embryonic, L4 and day 1 adult stages (**Fig. S2D**). We confirmed previously reported CFI-1 expression sites ^65,70^, but also identified new sites of expression in several neurons (CAN, RIP, SIB, REM) (**Table S1**, **Fig. S3I**). Altogether, our analysis established a nervous system-wide map of CFI-1 expression in *C. elegans*.

### *cfi-1* activity in PVC interneurons is required for posterior touch responses

Using the CFI-1 expression map, we looked for candidate neuron types within the *C. elegans* mechanosensory circuit (**Fig. 2C**). Notably, CFI-1 is not expressed in any mechanosensory neurons, suggesting it may act in interneurons and/or motor neurons (**Fig. 2C, Table S1**). Within the posterior touch circuit, *cfi-1* is expressed in two interneuron types (PVC and LUA) in the lumbar ganglion of the tail and two motor neuron types (DB, VB) along the ventral nerve cord (**Fig. 2C**). To determine whether *cfi-1* is required in DB and VB motor neurons ^73^, which mediate forward locomotion upon posterior touch, we assayed animals carrying a regulatory allele of *cfi-1* that specifically lacks expression in ventral nerve cord motor neurons, referred to as “*cfi-1 Motor Neuron-Knock Out (cfi-1 MN-KO)”* ^55^ (**Fig. 2A, D**). We found no significant differences between control and *cfi-1 MN-KO* animals (**Fig. 2E**), indicating that *cfi-1* activity in motor neurons is dispensable for posterior touch responses. Thus, we characterized interneurons next.

Previous studies using cell ablation, optogenetic, and calcium imaging have firmly established that PVC interneurons are essential for *C. elegans* posterior touch response ^59 74 67 75^, whereas LUA has been described as a “connector cell” between tail mechanosensory neurons and interneurons (**Fig. 2C**) ^59^. To determine whether *cfi-1* is required in LUA or PVC lumbar interneurons for normal posterior touch responses, we conducted cell-specific RNAi knockdown ^76^. LUA-specific RNAi against *cfi-1* showed no significant differences in the posterior harsh touch response (**Fig. 2F**), possibly due to incomplete *cfi-1* knockdown. In contrast, PVC-specific RNAi led to defects in posterior harsh touch response (**Fig. 2G**), albeit weaker than those observed in *cfi-1(ot786)* LOF mutants. Consistent with a recent report ^66^), our analysis demonstrates a cell-autonomous requirement for *cfi-1* in PVC interneurons for posterior harsh touch response.

### PVC and LUA lumbar interneurons are normally generated in *cfi-1* LOF mutants

Although PVC and LUA differ in neurotransmitter usage (PVC is cholinergic, LUA is glutamatergic), they are sister interneurons generated by the same mother cell during *C. elegans* embryogenesis (**Fig. S4A**). By mining protein and mRNA datasets of *C. elegans* embryogenesis ^72^ ^77^, we observed that *cfi-1* mRNA is first detected in the grandmother cell (ABplpppaap/ABprpppaap), whereas CFI-1 protein first appears in the mother cell (ABplpppaapa/ABprpppaapa) of PVC and LUA (**Fig. S4A**). Integration of larval scRNA-Seq datasets with our endogenous fluorescent reporter analyses shows that CFI-1 remains expressed in both PVC and LUA interneurons throughout development and adulthood (**Fig. 2B-C, S2D**)^78^. Given the early onset of *cfi-1* expression in the PVC/LUA lineage (**Fig. S4A**), we asked whether PVC and/or LUA interneurons are normally generated in *cfi-1* LOF mutants. We found this to be the case as the total number of tail neurons expressing a pan-neuronal reporter gene (*rab-3::yfp*) is unaffected in *cfi-1* LOF animals (**Fig. S2E**).

### CFI-1 controls the terminal identity of PVC and LUA lumbar interneurons

To determine the putative target genes of CFI-1 in PVC interneurons, we followed an unbiased approach. After integrating available ChIP-seq data for CFI-1^70^ with published expression profiles of PVC neurons at the L4 stage ^78^ (www.cengen.org), we found that CFI-1 binds to 526 (52.6%) of the top 1,000 highly expressed genes in PVC interneurons (**Fig. 3A, File S1**). Next, we employed TargetOrtho2, an *in silico* phylogenetic footprinting approach for the discovery of TF binding sites conserved across multiple nematode species ^79^. TargetOrtho2 predicted CFI-1 binding sites (motifs) in 395 (75.1%) of these 526 genes (**Fig. 3A, File S2**), suggesting that CFI-1 binds directly on DNA to control gene expression in PVC neurons.

**Figure 3.**
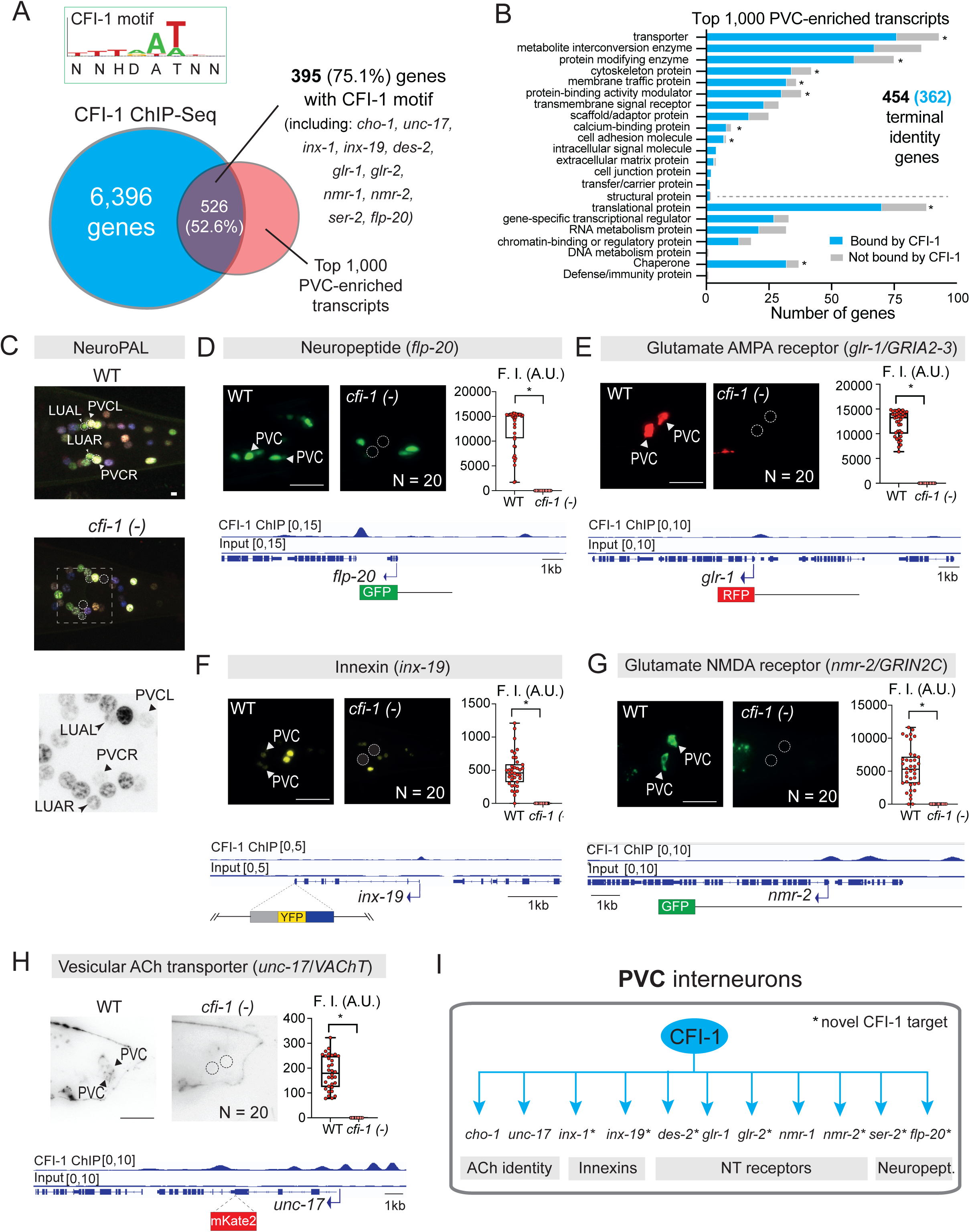
CFI-1 controls PVC interneuron identity. **A**. Top: DNA binding motif of CFI-1 ^70,129^. Bottom: Venn diagram of CFI-1-bound genes from ChIP-seq ^70^ and top 1,000 genes expressed in PVC ^78^. **B**. Protein class ontology analysis (Panther 19.0) of CFI-1 targets in PVC. *: significantly enriched categories (overrepresented relative to entire genome) *, p < 0.01. Fisher’s exact. **C**. Representative images of NeuroPAL strain in WT and *cfi-1(ot786)* adult (D1) animals. Inset with inverted colors is shown for better contrast. Dashed circles: LUA and PVC. Large arrowheads = PVC, small arrowheads = LUA. N = 10. **D-H.** Expression analysis of five PVC fluorescent reporters (*flp-20, glr-1, inx-19, nmr-2* and *unc-17*) in WT and *cfi-1(ot786)* adult (D1) animals. Top left: Representative images showing reporter expression. Arrowheads: PVC expression. Dashed circle: no PVC expression. Bottom: ChIP-Seq tracks for CFI-1 at its target genes and diagram of reporter used. Top right: Box and whisker plots displaying quantitative fluorescence intensity (F.I) measurements in arbitrary units (A.U.), with the whiskers extending to the minimum and maximum values, and the horizontal line within the box representing the median. Each dot represents an individual neuron. An unpaired *t*-test (two-sided) with Welch’s correction was performed: *, p < 0.0001 versus WT. Scale bars: 20 μm. **I.** Summarizing of CFI-1 targets in PVC. *: novel CFI-1 targets.

To gain insight into the categories of genes possibly regulated by CFI-1, we conducted gene ontology (GO) analysis on the top 1,000 PVC enriched genes (Panther 19.0) ^80^. We found an overrepresentation (454 genes, 45.4 %) of terminal identity genes (e.g., transporters, receptors, ion channels) (**Fig. 3B**), most of which (362 genes, 69%) are bound by CFI-1 (**Fig. 3B, File S3**).

We conducted a similar analysis for LUA interneurons. We found that CFI-1 binds to 643 (64.3%) of the top 1,000 LUA-enriched genes (**Fig. 4A, File S1**). TargetOrtho2 predicted CFI-1 binding sites (motifs) in 504 (78.6%) of these 643 genes (**Fig. 4A, File S2**). GO analysis also revealed an overrepresentation of terminal identity genes among the 1,000 LUA-enriched genes with 416 (41.6%) fitting in this category (**Fig. 4B**), most of which (289 genes, 72%) are bound by CFI-1 (**Fig. 4B, File S4**). Altogether, both ChIP-Seq and *in silico* prediction support the hypothesis that CFI-1 directly regulates the expression of PVC and LUA terminal identity genes.

**Figure 4.**
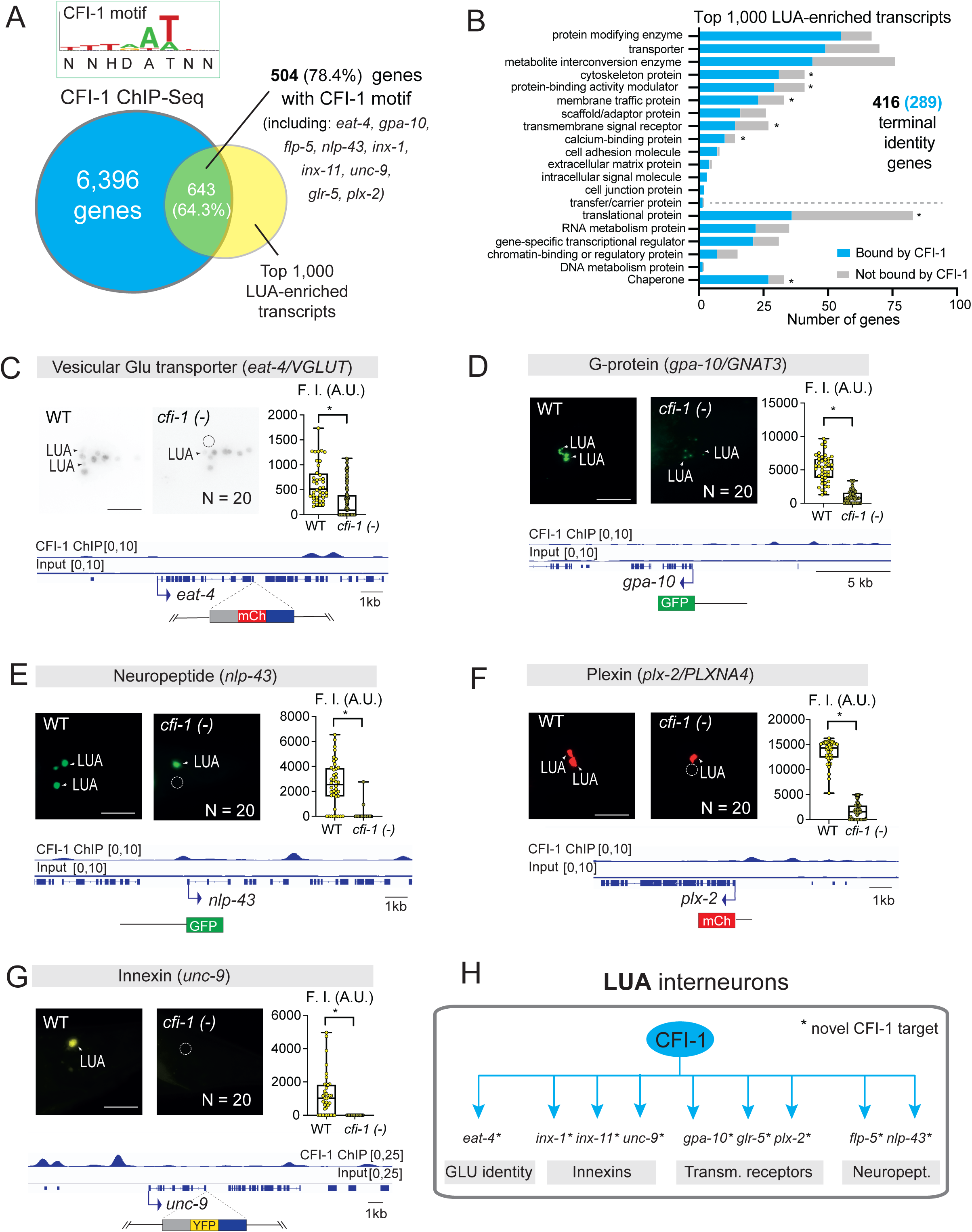
CFI-1 controls LUA interneuron identity. **A.** (Top) Directly determined binding motif of CFI-1 ^70,129^. (Bottom) Venn diagram summarizing the intersection of CFI-1-bound genes from ChIP-seq ^70^ and the top 1000 genes expressed in LUA interneurons from CenGEN ^78^. **B.** Graph summarizing protein class ontology analysis (Panther 19.0) of CFI-1 target genes in LUA interneurons. GO categories that are significantly enriched (overrepresented relative to the entire genome) have an asterisk, *, p < 0.01 based on Fisher’s exact test. **C-G.** Expression analysis of five fluorescent reporters of LUA terminal identity (*eat-4, gpa-10, nlp-43, plx-2* and *unc-9*) in WT and *cfi-1(ot786)* mutant young adult animals. (Top left) Representative images showing reporters expression. White arrowheads indicate expression in LUA interneurons, while the white dashed circle marks the absence of expression in LUA interneurons (Bottom) IGV snapshots showing CFI-1 binding at *loci* of genes and a diagram of the reporter allele used. (Top right) box and whisker plots displaying quantitative fluorescence intensity (F.I) measurements in arbitrary units (A.U.), with the whiskers extending to the minimum and maximum values, and the horizontal line within the box representing the median. Each dot represents an individual neuron. An unpaired *t*-test (two-sided) with Welch’s correction was performed: *, p < 0.0001; n.s., not significant versus WT. Scale bars: 20 μm. **H.** Diagram summarizing the gene targets of CFI-1 in LUA interneurons. Asterisks denote CFI-1 novel targets.

To test this hypothesis, we initially used the NeuroPAL multicolor reporter strain, as it can uncover changes in terminal identity caused by TF mutations ^71^. We observed a fluorescent color change in NeuroPAL signal specifically in PVC and LUA interneurons of *cfi-1* LOF mutants (**Fig. 3C, S5G, S6G**), indicative of terminal identity defects. To precisely determine which genes are affected, we obtained and independently validated dozens of available terminal identity markers for PVC and LUA interneurons (**File S5**). Next, we systematically tested their expression in WT and *cfi-1* LOF animals. In PVC interneurons, the expression of 11 out of the 13 tested reporters was either completely absent (*flp-20/FMRF-like peptide*, *glr-1/GRIA2-3*, *inx-19/innexin*, *nmr-2/GRIN2C*, *unc-17/VAChT*) (**Fig. 3D-H, Table S2**), or significantly reduced (*cho-1/ChT*, *des-2/* nAChR, *glr-2/GRIA1-3*, *nmr-1/GRIN1*, *ser-2*/GPCR, *inx-1*/innexin) in *cfi-1* mutants (**Fig. S5A-E, S6D, Table S2**). Four of these 11 genes (*unc-17/VAChT, cho-1/ChT, glr-1/GRIA2-3, nmr-1/GRIN1*) are independently confirmed CFI-1 targets in PVC neurons ^47,65^. Further, ChIP-Seq binding peaks for CFI-1 are present in each of these 11 genes (**Fig. 3D-H, S5A-E, Table S2**).

In LUA interneurons, a similar approach revealed that the expression of 9 out of the 11 tested reporters was significantly reduced in *cfi-1* mutants (*eat-4/VGluT1-3*, *glr-5/GRIK4, gpa-10/G protein, flp-5/ FMRF-like peptide, nlp-43/neuropeptide, plx-2/Plexin2, innexins: inx-1, inx-11, unc-9*) (**Fig. 4C-G, S6A-D, Table S3**). Moreover, ChIP-seq confirmed that CFI-1 binds to all nine genes (**Fig. 4C-G, S6A-D**). Altogether, these findings provide strong evidence that CFI-1 functions as a terminal selector in both PVC and LUA lumbar interneurons by activating directly the expression of distinct sets of terminal identity genes (**Fig. 3I, 4H**).

### CFI-1 functions cell-autonomously to control PVC and LUA terminal identity

To determine whether *cfi-1* functions cell-autonomously to control the terminal identities of PVC and LUA interneurons, we performed cell-specific RNAi knockdown. PVC-specific RNAi against *cfi-1* led to reduced expression of *glr-1/GRIA2-3* in PVC cells (**Fig. 5A**). Similarly, LUA-specific RNAi against *cfi-1* led to reduced expression of two terminal identity genes (*plx-2* and *gpa-10*) in LUA cells (**Fig. 5B-C**). Hence, CFI-1 operates cell-autonomously in PVC and LUA interneurons to control their terminal identity.

**Figure 5.**
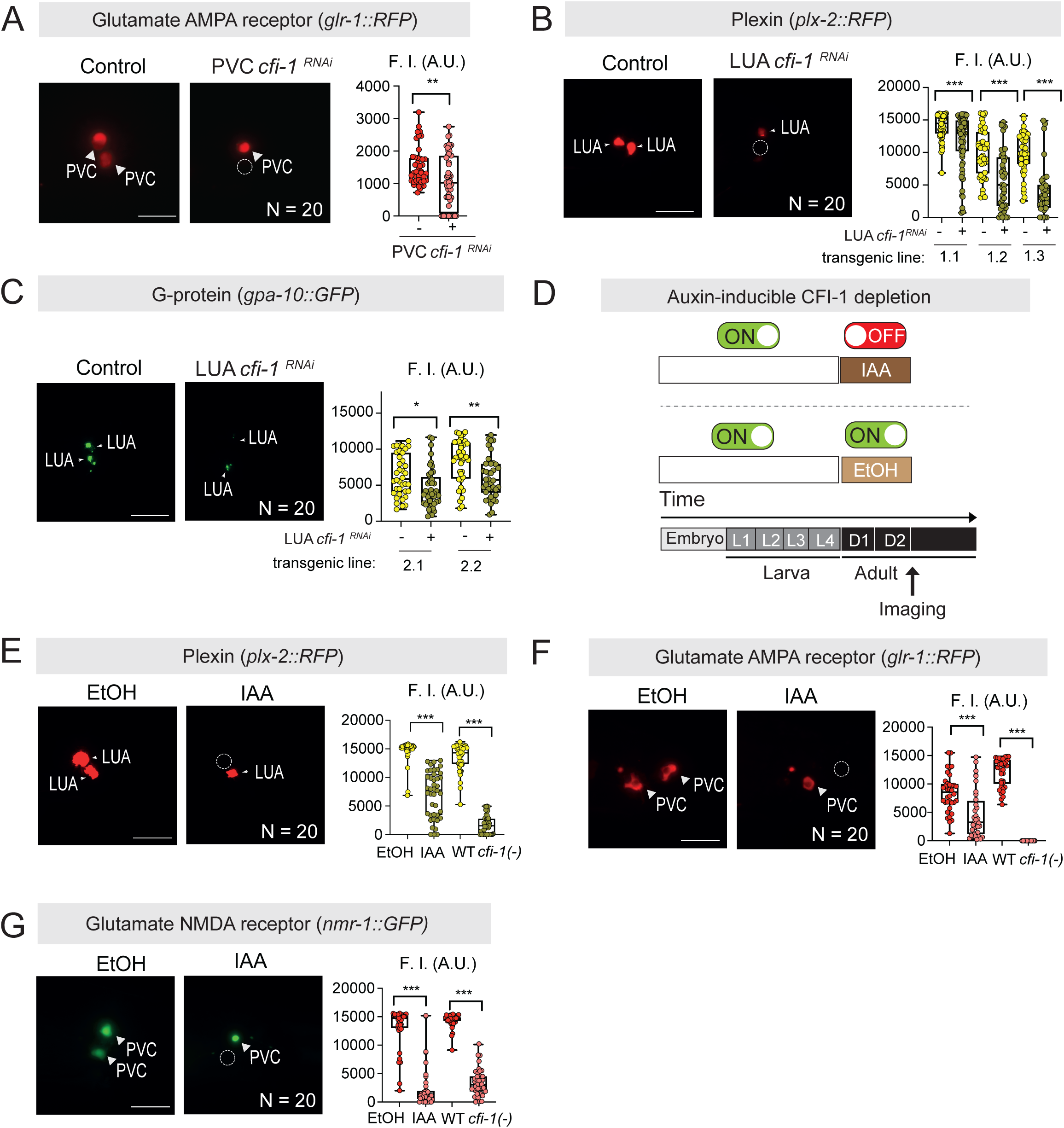
CFI-1 functions cell-autonomously to establish and maintain PVC and LUA interneuron identity. **A.** Representative images showing expression of *glr-1* reporter in the tail of control vs PVC-specific *cfi-1* RNA*i* in young adult animals. White arrowheads indicate expression in PVC interneurons, while white dashed circle indicates absence of expression in PVC interneuron. N = 20 animals. **B-C.** Representative images showing expression of *plx-2* or *gpa-10* reporters in the tail of control vs LUA-specific *cfi-1* RNA*i* in expressing young adult animals. White arrowheads indicate expression in LUA interneurons, while white dashed circles mark absence of expression in LUA interneurons. N = 20 animals. **D**. Diagrams illustrating the timeline of the ethanol (EtOH) control and auxin (IAA) treatments used for the depletion of CFI-1. Arrow indicates the time when fluorescence imaging was performed. **E**. Representative images showing expression of the *plx-2* reporter in the tail of control (EtOH) vs auxin (IAA) treated young adult animals. White arrowheads indicate expression in LUA interneurons, while white dashed circles mark absence of expression in LUA interneurons. N = 20 animals. **F-G.** Representative images showing expression of the *glr-1* or *nmr-1* reporter in the tail of control (EtOH) vs auxin (IAA) treated young adult animals. White arrowheads indicate expression in PVC interneurons, while white dashed circles mark absence of expression in PVC interneurons. N = 20 animals. In panel **D**, the green (ON) symbol represents presence of EGL-5 expression, while the red (OFF) symbol represents absence of EGL-5 expression. For panels **A**-**C** and **E-G**: Box and whisker plots displaying quantitative fluorescence intensity (F.I) measurements in arbitrary units (A.U.), with the whiskers extending to the minimum and maximum values, and the horizontal line within the box representing the median. Each dot represents an individual neuron. An unpaired *t*-test (two-sided) with Welch’s correction was performed: ^130^ *, p = 0.003; **, p < 0.002; ***, p < 0.0001, versus control (no RNAi); (**E**-**G**) ***, p < 0.0001 versus control (EtOH or WT). Scale bars: 20 μm.

### CFI-1 is required to maintain terminal identity features of PVC and LUA interneurons

Although our experiments with LOF alleles and cell-specific RNAi demonstrated that *cfi-1* is crucial for PVC and LUA terminal identities, they left unresolved whether CFI-1 is required in adulthood to maintain these identities. Using the AID system, we depleted CFI-1 in the first two days of adulthood (**Fig. 5D**). This led to a significant reduction in the expression of *plx-2* in LUA interneurons (**Fig. 5E**) and of *glr-1* and *nmr-1* in PVC interneurons after two days of auxin (IAA) treatment (**Fig. 5F-G**). Hence, CFI-1 is continuously required in early adulthood to maintain the expression of several terminal identity genes in both lumbar interneuron types (PVC and LUA).

### CFI-1 and EGL-5/HOX control the terminal identity of PVC and LUA interneurons

The incomplete penetrance of posterior touch defects (**Fig. 1B-C**) and residual expression of various PVC and LUA terminal identity genes in *cfi-1* mutants (**Fig. 4C-F, S5A-E, S6A-C**) suggest the involvement of additional TFs that partially compensate for the loss of *cfi-1*. The HOX cluster gene *egl-5,* a *C. elegans* homolog of posterior Abdominal-B-type HOX genes ^81^, emerged as a strong candidate TF for two reasons: First, *egl-5,* like *cfi-1*, is specifically required for posterior, but not anterior, touch responses ^82,83^. Second, within the posterior touch circuit, *egl-5* is expressed in mechanosensory (PLM) neurons, as well as in PVC and LUA interneurons ^84,85^. Since *cfi-1* and *egl-5* are co-expressed in PVC and LUA (**Fig. 6A, S4**), we hypothesized that they work together to control the terminal identity program of these cells.

**Figure 6.**
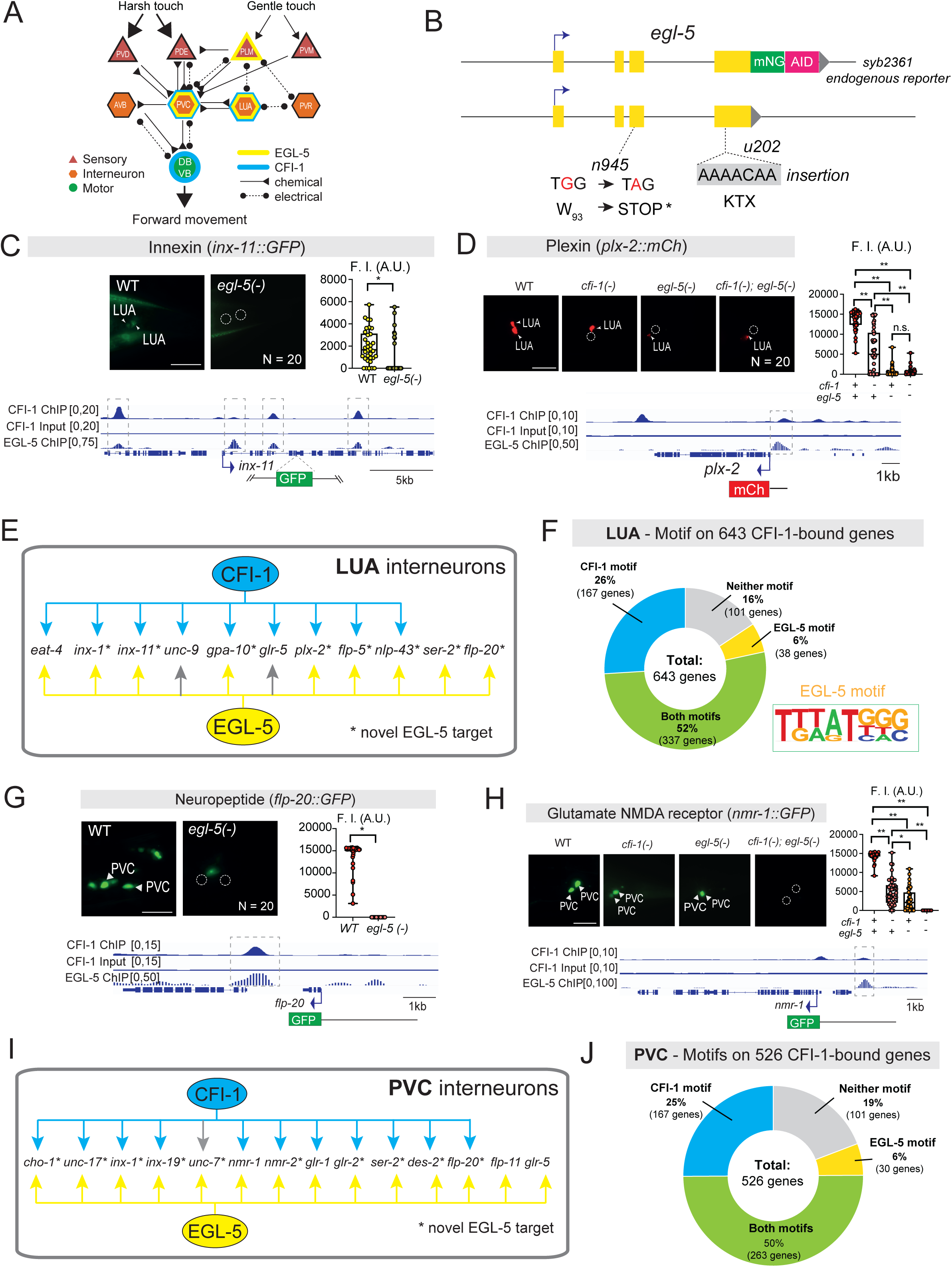
CFI-1 and EGL-5 jointly control PVC and LUA interneuron identity. **A.** Schematic illustrating the posterior touch circuit in *C. elegans*, highlighting CFI-1 (cyan outline) and EGL-5 (yellow outline) expression. Their expression only overlaps in LUA and PVC interneurons. **B.** Diagram of the *egl-5* locus. *n945* allele: G-to-A mutation in the third exon generates a premature STOP ^87^; *u202* allele: a insertion AAAACA in the fourth exon generates a frameshift variant ^86^; *syb2361* allele: *egl-5* endogenous reporter with an in-frame fluorescent protein mNeonGreen (mNG) insertion followed by a degron sequence immediately before the stop codon of *egl-5*. **C.** Expression analysis of an *inx-11* fluorescent reporter for LUA terminal identity in WT and *egl-5(n945)* mutant young adult animals. N = 20 animals. **D.** Expression analysis of a *plx-2* fluorescent reporter of LUA terminal identity in WT, *cfi-1(ot786), egl-5(n945)* single mutants and *cfi-1(ot786); egl-5(n945)* double-mutant young adult animals. N = 20 animals. **E.** Diagram summarizing the gene targets of CFI-1 and EGL-5 in LUA interneurons. Asteriks denote ELG-5 novel targets. Gray arrows: genes not tested. **F.** Donut chart summarizing the presence of CFI-1 and/or EGL-5 DNA-binding motifs on the CFI-1 bound genes in LUA interneurons. de novo motif discovery analysis of EGL-5 binding peaks identifies an 8 bp-long EGL-5 binding motif. **G.** *flp-20* fluorescent reporter of PVC terminal identity in WT and *egl-5(n945)* mutant young adult animals. N = 20 animals. **H.** Expression analysis of a *nmr-1* fluorescent reporter of PVC terminal identity in WT, *cfi-1(ot786), egl-5(n945)* single mutants and *cfi-1(ot786); egl-5(n945)* double-mutant young adult animals. N = 20 animals. **I.** Diagram summarizing the gene targets of CFI-1 and EGL-5 in PVC interneurons. Asteriks denote ELG-5 novel targets. Gray arrow: gene that was not tested. **J.** Donut chart summarizing the presence of CFI-1 and/or EGL-5 DNA-binding motifs on the CFI-1 bound genes in PVC interneurons. For panels **C**-**D** and **G-H**: (Top left) Representative images showing reporters expression. WT image shown is the same in both cfi-1 and egl-5 mutant analyses. White arrowheads indicate expression in LUA or PVC interneurons, while the white dashed circle marks the absence of expression in LUA or PVC interneurons. (Bottom) IGV snapshots showing CFI-1 and EGL-5 binding at *loci* of genes and a diagram of the reporter allele used. EGL-5 input track is not shown because original authors normalized to the EGL-5 ChIP signal. (Top right) Box and whisker plots displaying quantitative fluorescence intensity (F.I) measurements in arbitrary units (A.U.), with the whiskers extending to the minimum and maximum values, and the horizontal line within the box representing the median. Each dot represents an individual neuron. An unpaired *t*-test (two-sided) with Welch’s correction was performed: (**C**, **G**) *, p < 0.0001 versus WT; (**D, H**) *, p = 0.0001; **, p < 0.0001; n.s., not significant versus single or double mutant. Scale bars: 20 μm.

To test this, we first examined the role of *egl-5* in LUA neurons using a strong LOF *egl-5 (n945)* allele, for which Sanger sequencing revealed a premature STOP codon in exon 3 (**Fig. 6B**) ^86^ ^87^. We tested nine LUA terminal identity markers (GLU biosynthesis: *eat-4/VGLUT;* neuropeptides: *flp-5, flp-20, nlp-43;* innexin: *inx-1, inx-11;* receptors: *gpa-10, ser-2, plx-2,*) and all of them were affected in *egl-5 (n945)* animals (**Fig. 6C-D, S8, Table S3**). Prior work showed that *eat-4/VGLUT* is also affected in LUA neurons of animals carrying another *egl-5(u202)* allele (**Fig. 6B**) ^88^. Notably, the LUA expression of seven of these nine genes also depends on *cfi-1* (**Fig. 6E**, **Table S3**), suggesting that EGL-5 and CFI-1 share transcriptional target genes in LUA interneurons. Further, ChIP-seq analysis showed that EGL-5 and CFI-1 bind at the *cis*-regulatory regions of all tested LUA terminal identity genes (**Fig. 6C-D, S8, Table S3**). Consistently, TargetOrtho2 predicted both EGL-5 and CFI-1 binding sites in all of these 7 genes (**Fig. S10J-O, File S2**), and more broadly in 337 of the 643 (52%) genes bound by CFI-1 (ChIP-Seq) (**Fig. 6F**). The EGL-5 and CFI-1 sites on their shared target genes are separated by dozens or hundreds of nucleotides (**Fig. S10D-O), File S2**), arguing against a cooperative model where EGL-5 and CFI-1 bind DNA as a heterodimer.

With a similar analysis, we investigated the function of *egl-5* in PVC interneurons. We identified 10 terminal identity genes (NT biosynthesis: *unc-17/VAChT, cho-1ChT;* neuropeptide: *flp-20;* NT receptors: *glr-2, nmr-2, ser-2, des-2;* innexins: *inx-1, inx-19, unc-7*) as new EGL-5 targets in PVC interneurons (**Fig. 6G, S7, S8G**). Prior work showed that *egl-5* controls the expression of four additional terminal identity genes (*glr-1, nmr-1, glr-5, flp-11*) ^83,85^. Notably, 11 out of these 14 PVC-specific terminal identity genes are also affected in *cfi-1* LOF mutants (**Fig. 6I**, **Table S2**), suggesting that CFI-1 and EGL-5 share transcriptional targets in PVC interneurons as well. Supporting the notion of TF collaboration, we observed stronger effects on *nmr-1* expression in *cfi-1; egl-5* double LOF mutants compared to either single mutant (**Fig. 6H**). Further, ChIP-Seq showed that both EGL-5 and CFI-1 bind to the *cis*-regulatory region of 13 of these 14 genes (**Fig. 6G-H, S7, S8G Table S2**). Importantly, TargetOrtho2 predicted both EGL-5 and CFI-1 binding sites in all 13 genes, which were again physically separated by dozens of nucleotides (**Fig. S10D-K, File S2**). Sites for both TFs were also found in 263 of the 526 (50%) genes bound by CFI-1 (ChIP-Seq) (**Fig. 6J**). Altogether, we conclude that CFI-1 and the posterior Hox protein EGL-5 cooperate to directly activate the expression of distinct sets of terminal identity genes in LUA and PVC interneurons (**Fig. 6E, I**).

**Figure 7.**
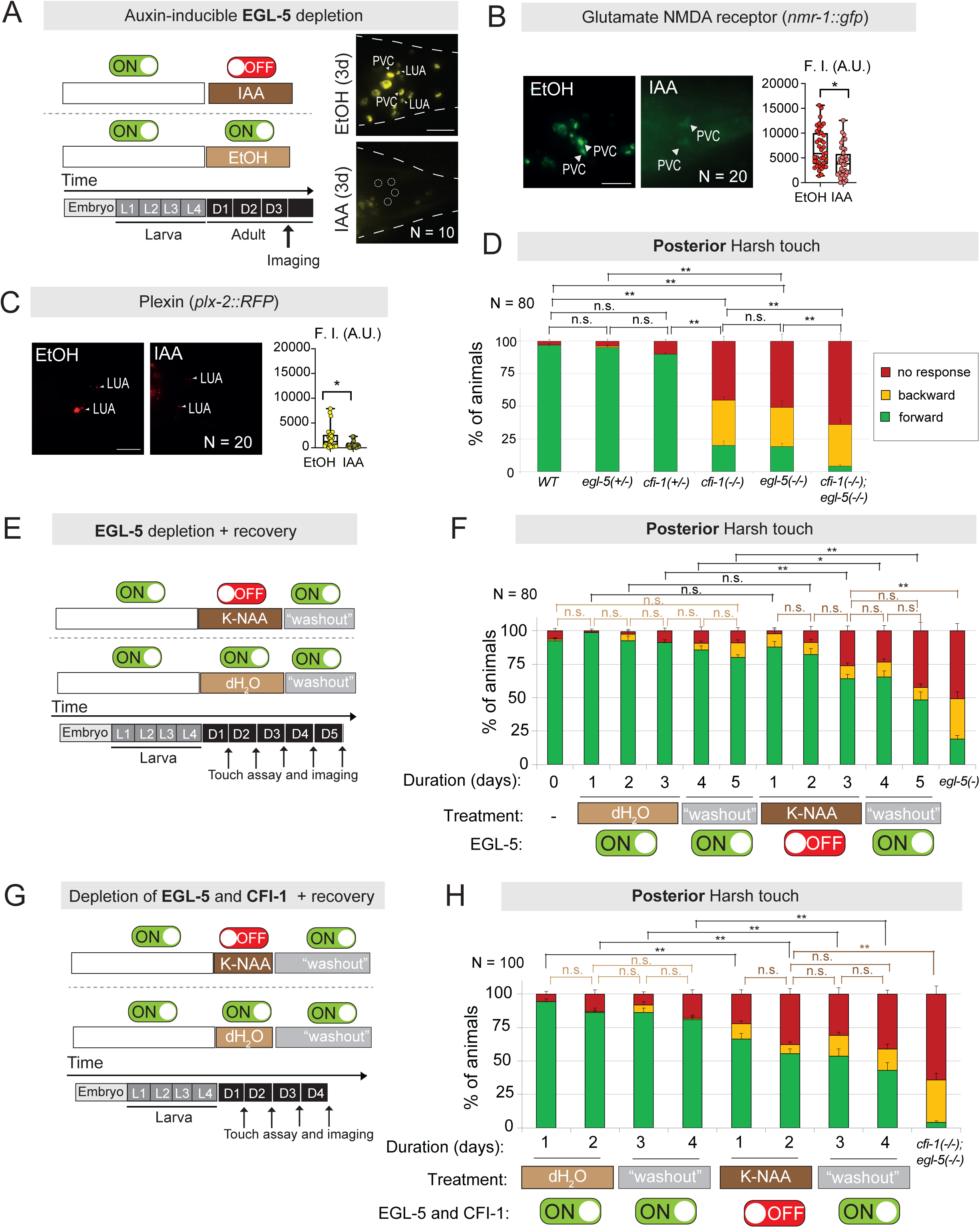
EGL-5 is required to maintain interneuron terminal identity and normal posterior touch responses. **A.** Diagram illustrating the timeline of the EtOH (control) and auxin (IAA) treatments used for the depletion of EGL-5. Arrow indicates the time when fluorescence imaging was performed. Representative images showing EGL-5::mNG::AID expression in the tail of *C*. *elegans* animals after 3 days of control (EtOH) and auxin (IAA) treatments. N = 10 animals. **B.** Representative images showing expression of the *nmr-1* reporter in the tail of EtOH (control) vs auxin (IAA) treated young adult animals. White arrowheads indicate expression in PVC interneurons. Graph showing fluorescence intensity quantifications of the *nmr-1* reporter in PVC interneurons. N = 20 animals. **C.** Representative images showing expression of the *plx-2* reporter in the tail of EtOH (control) vs auxin (IAA) treated young adult animals. White arrowheads indicate expression in LUA interneurons. Graph showing fluorescence intensity quantifications of the *plx-2* reporter in LUA interneurons. N = 20 animals. **D.** Posterior harsh touch assay on *C. elegans* animals carrying *cfi-1* and *egl-5* mutations, either alone or in combination, as well as heterozygous animals for each allele. N = 80 animals. **E.** Diagrams illustrating the timeline of the dH2O (control) and auxin (K-NAA) treatments used for the depletion and recovery of EGL-5. Arrows indicate when touch assays and fluorescence imaging were performed. **F.** Posterior harsh touch assay on *C. elegans* animals with no treatment, and after dH2O (control), auxin (K-NAA) and “washout” recovery treatments along with *egl-5(n945)* mutant animals for comparison. N = 80 animals. **G.** Diagrams illustrating the timeline of the dH2O (control) and auxin (K-NAA) treatments used for the depletion and recovery of both CFI-1 and EGL-5. Arrows indicate when touch assays and fluorescence imaging were performed. **H.** Posterior harsh touch assay on *C. elegans* animals with no treatment, and after dH2O (control), auxin (K-NAA) and “washout” recovery treatments along with *cfi-1(ot786)*; *egl-5(n945)* double-mutant animals for comparison. N = 100 animals. In panels **A**, **E-F**, the green (ON) symbol represents presence of EGL-5 expression, and the red (OFF) symbol represents absence of EGL-5 expression. While in panels **G-H**, the green (ON) symbol represents presence of both CFI-1 and EGL-5 expression, and the red (OFF) symbol represents absence of both CFI-1 and EGL-5 expression. For panels **B-C**: Box and whisker plots displaying quantitative fluorescence intensity (F.I) measurements in arbitrary units (A.U.), with the whiskers extending to the minimum and maximum values, and the horizontal line within the box representing the median. Each dot represents an individual neuron. An unpaired *t*-test (two-sided) with Welch’s correction was performed: (**B**) *, p = 0.0033, versus control (EtOH); (**C**) *, p = 0.0006, versus control (EtOH). Scale bars: 20 μm. For behavior in panels **D**, **F** and **H**: *, p = 0.0056; **, p < 0.0001; n.s., not significant, as determined by Bonferroni post-hoc tests.

### EGL-5 is required to maintain terminal identity features of PVC and LUA interneurons

By mining protein and mRNA datasets of *C. elegans* embryogenesis ^72,77^, we found that *egl-5*, like *cfi-1*, is first detected in the grandmother cell (ABplpppaap/ABprpppaap) of PVC and LUA interneurons (**Fig. S4A-B**). Further, analysis of our endogenous *egl-5::mNG::AID* reporter allele revealed continuous expression of EGL-5/Hox in PVC and LUA cells during larval and adult stages (**Fig. 6A, 7A, S4D**). The persistent expression of a Hox protein in adult neurons is unusual, as Hox genes are well known for their early patterning roles^28^. To determine whether EGL-5 is required in adult neurons to maintain the expression of PVC and/or LUA terminal identity genes, we again employed the AID system. By using our auxin-inducible *egl-5::mNG::AID* allele (**Fig. 6B**), we exposed L4 animals to natural auxin (IAA) for 3 days and confirmed robust EGL-5::mNG::AID depletion in PVC and LUA interneurons, as well as other tail cells (**Fig. 7A**). Adult-specific depletion of EGL-5 led to a significant decrease of *nmr-1* in PVC interneurons (**Fig. 7B**) and *plx-2* in LUA interneurons (**Fig. 7C**). We conclude that EGL-5 is continuously required to sustain the expression of terminal identity genes both in PVC and LUA interneurons, uncovering a non-canonical role for a Hox patterning gene in maintaining adult neuron identity.

### EGL-5 cooperates with CFI-1 to control posterior touch responses

We complemented our terminal identity marker analysis with behavioral assays on animals carrying strong LOF alleles for *egl-5 (n945)* or *cfi-1 (ot786).* We found that *egl-5 (n945)* and *cfi-1 (ot786)* homozygous mutants exhibit strikingly similar posterior harsh touch phenotypes (**Fig. 7D**). No defects were observed in heterozygous *egl-5 (n945/+)* and *cfi-1 (ot786/+)* animals, indicating that a single copy of either gene is sufficient for normal posterior touch responses (**Fig. 7D**). Notably, *cfi-1 (ot786)*; *egl-5 (n945)* double mutants displayed more severe posterior touch defects compared to either single mutant (**Fig. 7D**), suggesting that *cfi-1* and *egl-5* cooperate to control PVC and LUA identity, and thereby touch circuit function.

### EGL-5 is required in the adult stage to control posterior touch behavior

Since EGL-5 is required to maintain adult PVC and LUA terminal identity (**Fig. 7A-C**), we wondered whether it is also necessary to maintain normal touch-evoked behavioral responses. Using the AID system, we exposed L4 animals to synthetic auxin (K-NAA) for 3 consecutive days (**Fig. 7E**). Following the depletion of EGL-5::mNG::AID in PVC and LUA interneurons (**Fig. 7F, S9A-D**), we observed a significant impairment in posterior harsh touch responses at 3 days (D3) after K-NAA treatment initiation, but not earlier (**Fig. 7F**). Importantly, a washout period of two days (D4 and D5) did restore EGL-5::mNG::AID expression (**Fig. S9A-D**), but such restoration was insufficient to improve posterior harsh touch defects (**Fig. 7F**). We conclude that EGL-5, like CFI-1, is required to maintain normal posterior touch responses, implicating a Hox patterning gene in the maintenance of adult circuit function.

In contrast to CFI-1 though, EGL-5 does not function as a reversible on-off switch, i.e., following its depletion in adult PVC and LUA cells, EGL-5 resupply did not restore posterior touch responses (**Fig. 7F**). Intriguingly, we found that simultaneous depletion of both CFI-1 and EGL-5 with the AID system in early adulthood (D1-2) led to posterior harsh defects, which were not reversible following a “wash-out” period of two days (D3-4) (**Fig. 7G-H, S9E-H**). Hence, the unique ability of CFI-1 to restore adult circuit function depends on CFI-1.

### CFI-1 and EGL-5 operate in a double-positive feedback loop in PVC and LUA interneurons

Our findings indicate that CFI-1 and EGL-5 act as terminal selectors both in PVC and LUA interneurons by activating distinct sets of terminal identity genes (**Fig. 6 E, I**). Because ChIP-Seq revealed extensive binding of EGL-5 to the *cis*-regulatory region of *cfi-1* and vice versa (**Fig. 8A, D**), we investigated a potential cross-regulatory relationship. Indeed, expression of the endogenous mNG::AID::CFI-1 reporter is significantly reduced both in PVC and LUA interneurons of *egl-5* LOF mutants (**Fig. 8A**). Similarly, expression of an endogenous EGL-5::mNG::AID reporter is diminished in PVC and LUA neurons of *cfi-1* LOF mutants (**Fig. 8D**). Hence, CFI-1 and EGL-5 are mutually required to activate each other’s expression, and jointly required to activate distinct terminal identity genes in PVC and LUA interneurons. These findings unveil a double-positive feedback loop that “locks-in” PVC and LUA terminal identity (**Fig. 8G-H**).

**Figure 8.**
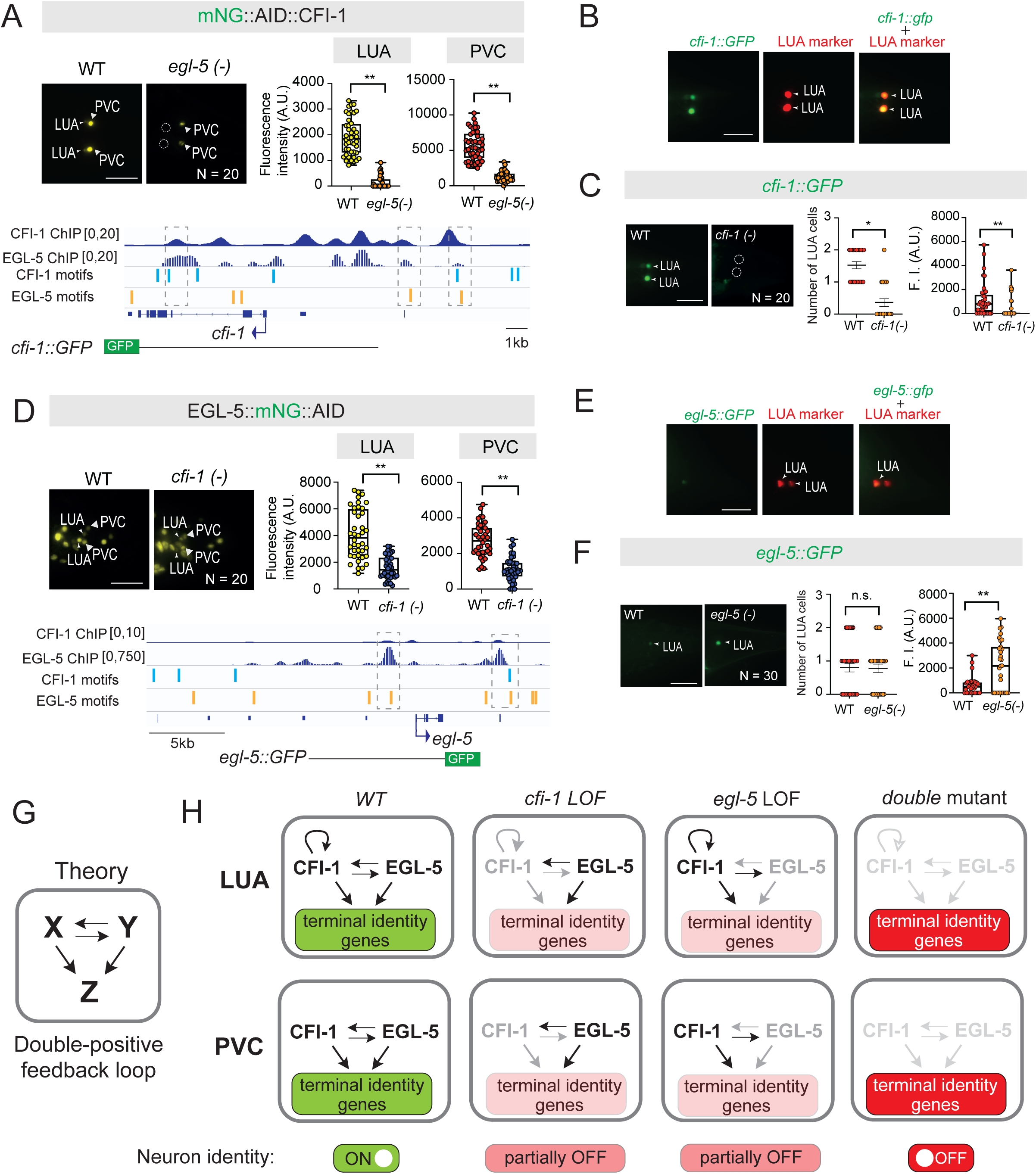
A double-positive feedback loop ensures robustness of PVC and LUA interneuron identity. **A.** Expression analysis of a CFI-1 fluorescent reporter in WT and *egl-5(n945)* mutant young adult animals. N = 20 animals. **B.** Representative images of the worm tail region showing CFI-1::GFP expression in combination with a LUA reporter (*plx-2::mch*). White arrowheads indicate expression in LUA interneurons. N = 10 animals. **C.** Expression analysis of CFI-1::GFP reporter in WT and *cfi-1(ot786)* mutant young adult animals, N = 20 animals. **D.** Expression analysis of an EGL-5 fluorescent reporter in WT and *cfi-1(ot786)* mutant young adult animals. N = 20 animals. **E.** Representative images of the worm tail region showing EGL-5::GFP expression in combination with a LUA reporter (*plx-2::mch*). White arrowheads indicate expression in LUA interneurons. N = 10 animals. **F.** Expression analysis of EGL-5::GFP reporter in WT and *egl-5(n945)* mutant young adult animals, N = 20 animals **G.** Theoretical model summarizing the double-positive feedback loop. **H.** Schematic model summarizing the double-positive feedback loop regulating the identity of LUA and PVC interneurons, and the CFI-1 autoregulation in LUA interneurons. For panels **A** and **D**: (Top left) Representative images displaying reporter expression. White arrowheads indicate expression in LUA and/or PVC interneurons, while the white dashed circles mark the absence of expression in LUA interneurons. (Bottom) IGV snapshots showing CFI-1 and EGL-5 binding at *cfi-1* or *egl-5* gene locus, together with their DB motifs found by TargetOrtho 2 analysis and a diagram of the reporter alleles showed in panels **B-C** and **E-F**. Dashed boxes highlight the CFI-1 and/or EGL-5 DB motifs that are in the same region of the ChIP peaks. (Right) Box and whisker plots displaying quantitative fluorescence intensity (F.I) measurements in arbitrary units (A.U.), with the whiskers extending to the minimum and maximum values, and the horizontal line within the box representing the median. Each dot represents an individual LUA or PVC neuron. For panels **C** and **F,** (Left graph) Scatter dot plots displaying number of LUA neurons expressing respective fluorescent reporter, with the horizontal line representing the mean with standard error of the mean (SEM). Each dot represents the average number of reporter (+) LUA neurons in one animal. (Right graph) Box and whisker plots displaying quantitative fluorescence intensity (F.I) measurements in arbitrary units (A.U.), with the whiskers extending to the minimum and maximum values, and the horizontal line within the box representing the median. Each dot represents an individual neuron. An unpaired *t*-test (two-sided) with Welch’s correction was performed: *, p = 0.0011; **, p < 0.0001 versus WT. Scale bars: 20 μm.

### CFI-1 positively regulates its own expression in LUA interneurons

How is *cfi-1* and *egl-5* expression maintained in adult PVC and LUA interneurons? ChIP-Seq indicates that CFI-1 and EGL-5 bind to their own loci (**Fig. 8A, D**), raising the possibility of direct transcriptional autoregulation. To test this, we first characterized available *cfi-1* and *egl-5* transgenic reporters ^55,65,89^. We found no expression in PVC cells for both *cfi-1* and *egl-5* reporters; it is likely that PVC-specific *cis*-regulatory elements are missing from these transgenic reporters (**Fig. 8A, D**). However, we identified two *cfi-1* reporters with specific expression in LUA interneurons (**Fig. 8A-B, S10A-C**), enabling us to test whether CFI-1 autoregulates in these cells. Indeed, the expression of both reporters is significantly reduced in *cfi-1* LOF mutants (**Fig. 8C, S10B**). Hence, we favor a model where *cfi-1* binds to its own locus and positively autoregulates to perpetuate its expression in LUA interneurons (**Fig. 8H**). Intriguingly, this is not the case for *egl-5*. Expression of an *egl-5* reporter is not reduced in LUA interneurons of *egl-5* LOF mutants (**Fig. 8D-F**). Altogether, we propose that positive CFI-1 autoregulation reinforces and actively stabilizes the double-positive feedback loop in LUA interneurons (**Fig. 8G-H**). Autoregulation ensures sufficient CFI-1 levels to not only maintain *egl-5* but also the expression of both *cfi-1-* and *egl-5*-dependent genes, robustly securing interneuron identity and touch circuit function.

## DISCUSSION

In electrical engineering, an on-off switch is used to ensure circuit bistability, and thereby maintenance of two distinct states (ON and OFF) over time. By analogy, our findings in adult *C. elegans* uncover a functional “on-off” switch for a touch-evoked escape response circuit. Toggling between normal and low CFI-1/ARID3 levels, and vice versa, generated digital-like (ON/OFF) effects both on the molecular identity program of two lumbar interneuron types (PVC and LUA) and on animal behavior. Strikingly, resupply of CFI-1 effectively restored adult-onset defects both on interneuron identity and escape response. CFI-1 does not act alone, but it collaborates with a posterior HOX protein, EGL-5, to induce during development, and maintain in adulthood, a broad spectrum of PVC and LUA terminal identity features. Mechanistically, we identified two interconnected network motifs, a double-positive CFI-1/EGL-5 feedback loop and CFI-1 positive autoregulation, which “lock-in” the interneuron identity programs (**Fig. 8H**), and thereby secure information processing within the *C. elegans* posterior touch circuit. We discuss below the implications of our findings for the fields of developmental neurobiology, neurological disease, and systems neuroscience.

### Implications for terminal selector biology and developmental neuroscience

To understand how neuronal circuit function is preserved throughout life, it is essential to elucidate the molecular mechanisms that establish and maintain neuronal identity. The concept of terminal selectors aims to bridge this fundamental knowledge gap. Several studies in invertebrates and vertebrates support a model where a terminal selector, or combination thereof, controls the identity and function of a specific neuron type^33–35^ ^36–38^. However, our mechanistic understanding of how terminal selectors maintain neuronal identity remains poor, despite accumulating evidence linking human orthologs of terminal selectors to various neurodevelopmental and neurodegenerative disorders^39–46^. Our findings advance our understanding of neuronal identity control mechanisms in four major ways.

First, most terminal selectors to date have been described in either sensory or motor neuron types of worms, flies and mice ^31,39,56^. Here, we expand the terminal selector logic to interneurons of the *C. elegans* posterior touch circuit. By integrating genetic, biochemical (ChIP-Seq), and behavioral analyses, we demonstrate that CFI-1/ARID3 and EGL-5/HOX act as *bona fide* terminal selectors of PVC and LUA interneuron identity. By analyzing the expression of dozens of molecular markers, we unveil that CFI-1 and EGL-5 act directly to control a broad spectrum of terminal identity features ranging from NT biosynthesis to electrical synapses and neuropeptides, offering a blueprint for future comprehensive studies on terminal selector genes.

Second, accumulating evidence indicates that terminal selectors can act in combinations in a given neuron type^21,47,48,90,91^ ^92, 93,94^, highlighting the complexity of the transcriptional networks that control neuronal terminal identity. To understand their structure and function, it is important to identify network motifs - specific, recurring patterns of TF interactions and the genes they regulate. To date, the identification of such motifs remains a challenge for most terminal selectors with two notable exceptions. In *C. elegans*, CHE-1 operates within a bistable feedback loop of TFs and microRNAs to control chemosensory (ASE) neuron identity^91^. In *D. melanogaster*, Orthodenticle (Otd) and Defective proventriculus (Dve) operate in interlocked feedforward loops to control photoreceptor identity^92^. Here, we uncovered a different mechanism where CFI-1/ARID3 and EGL-5/HOX operate within a double-positive feedback loop to lock-in the terminal identity program of PVC and LUA interneurons (**Fig. 8H**). Double-positive feedback loops remain poorly understood ^95^, as opposed to the extensively studied double-negative feedback loops ^96,97^. We offer the first example of a double-positive TF feedback loop in an adult animal cell type, contributing to the systems biology field as most networks motifs have been primarily studied in unicellular systems ^95,98^, or during early animal development^99^.

Third, human genetic studies have linked several terminal selectors (e.g., *FEV, NURR1, EBF3*) to both neurodevelopmental and neurodegenerative conditions ^57^ ^40,43,44^. By leveraging the reversibility of the AID system, we show here that adult-onset loss of CFI-1 compromises interneuron identity and animal behavior. Strikingly, reintroduction of CFI-1 following its prolonged depletion restored escape response defects in the adult, a finding with biomedical implications for the restorability of neuronal and behavioral defects caused by mutations or variation in TF-encoding genes, which have been abundantly reported in the literature^100^ ^39, 101^. Notably, the gradual recovery of posterior harsh touch defects following adult-specific depletion and resupply of CFI-1 suggests that terminal selector depletion for an extended period (e.g., 2 days correspond to 15-20% of the adult *C. elegans* lifespan) does not permanently impair interneuron identity or circuit function. Instead, PVC and LUA identity is temporarily compromised, but these cells remain in a “responsive state” and thus capable of regaining their original terminal identity upon CFI-1 resupply, leading to restoration of circuit function. We emphasize that the reversible “on-off” switch function of CFI-1 is a result of experimental manipulations, i.e., CFI-1 depletion and resupply. Hence, it is fundamentally different from reversible switches that occur naturally, such as the *lactose (lac)* operon in *E. coli*^102^. Altogether, our findings suggest that terminal selectors are more versatile than previously thought; they are not only required to maintain adult neuron identity, but can also function as reversible genetic switches capable of effectively restoring adult neuron function.

Fourth, Hox genes are well-known for their fundamental roles in early nervous system patterning (e.g., cell specification, survival, and migration) ^28,62,103–105^. However, their putative neuronal functions during adult life remain largely unexplored. Here, we show that the posterior Hox gene *egl-5* (Abd-B/Hox9-13) is required – in the adult – to maintain interneuron terminal identity (PVC and LUA) and normal touch responses. Consistently, *egl-5* is known to induce the expression of terminal identity genes in mechanoreceptors (PLM) and other *C. elegans* tail neurons ^106^ ^83,85,88,107^, raising the possibility that it functions as a terminal selector in other neuron types. Of note, a previous study used a weaker *egl-5* allele *(u202)* (**Fig. 6B**), and concluded that *egl-5* is dispensable for LUA differentiation ^85,88^, contrasting our findings with strong LOF allele and protein depletion strategies. Further, the midbody Hox gene *lin-39* (Scr/Dfd/Hox4-5) is continuously required to maintain the cholinergic identity of adult *C. elegans* motor neurons ^108^. Altogether, these findings support a non-canonical role for Hox proteins as terminal selectors of neuronal identity.

### Two interlocked network motifs ensure robustness of touch interneuron identity

Computational modeling and studies in unicellular systems suggest that a double-positive feedback loop confers robustness, by locking a system into a steady state. Such loops are characterized by “joint bistability” ^95,109^ ^110^, where both factors (X and Y) are either both ON or OFF (**Fig. 8G**). An initial and transient input signal can drive the system to “lock-in” one of these steady states, ON or OFF. Although the initial signals that induce CFI-1 and EGL-5 in the PVC/LUA developmental lineage remain unknown, our findings suggest that CFI-1 (X) and EGL-5 (Y) operate in a double-positive feedback loop to “lock-in” the terminal identity program of LUA and PVC interneurons. We surmise that this loop robustly insulates the LUA and PVC identity program from potential fluctuations in *cfi-1* or *egl-5* expression. This is supported by our observations in heterozygous animals, where reduction of *cfi-1* or *egl-5* gene dosage by half had no apparent effect on posterior touch responses (**Fig. 7D**). However, homozygous loss of *egl-5* or *cfi-1* did lead to partial penetrant defects in ∼75% of the animals (**Fig. 7D**). Consistently, some terminal identity genes are strongly affected by *egl-5* or *cfi-1* loss (**Fig. 3-4, 6**), whereas others are only partially affected (**Fig. S5-8**). According to the systems biology definition of a double-positive feedback loop^97^, loss of either TF should result in the OFF state, and as such in fully penetrant defects. This discrepancy can be explained by the contribution of other, yet-to-be identified TFs that influence CFI-1 and EGL-5, or by the stabilizing effect of CFI-1 positive autoregulation. For example, removal of *egl-5* does not result in the OFF state because only the *egl-5* input onto *cfi-1* is affected, whereas the autoregulatory input is preserved (**Fig. 8H**). On the other hand, double LOF mutant analysis, which removes *egl-5* and *cfi-1* activity, as well as *cfi-1* autoregulation, likely leads the lumbar interneuron identity program to reach its OFF state as evident by the nearly fully penetrant defects on posterior touch responses (**Fig. 7H, 8G**) and terminal marker expression (**Fig. 6H**). Altogether, we propose that the double-positive feedback loop locks-in the terminal identity program of PVC and LUA interneurons. Self-perpetuation of CFI-1 expression by autoregulation further stabilizes this loop, ensuring the robustness of interneuron identity and touch-evoked escape responses.

### Why only CFI-1 is able to restore adult-onset defects in interneuron function?

The most parsimonious explanation is the dual input on CFI-1 (by itself and EGL-5), whereas EGL-5 receives a single input (by CFI-1) (**Fig. 8H**). Upon adult-specific CFI-1 depletion, a severe perturbation that impacts animal behavior, the combination of two activating inputs help CFI-1 regain a minimal expression threshold necessary for the re-establishment of interneuron terminal identity, which effectively restores circuit function. This model is supported by the key observation that combined depletion of both CFI-1 and EGL-5 in the adult resulted in failure to restore posterior touch defects (**Fig. 7G-H**). Further, positive autoregulation has been proposed as a mechanism to amplify the initial expression (induced by transient developmental signals) of a terminal selector to reach a minimal threshold required to initiate a specific neuron identity program ^111^. However, if a terminal selector solely requires itself to maintain its expression during adult life, then perturbations (e.g., noise, fluctuations) of terminal selector expression for extended periods of time could bring its expression below the needed threshold, resulting in catastrophic effects on neuron identity maintenance ^112,113^. Therefore, positive autoregulation is insufficient by itself, and additional mechanisms must be operating to ensure terminal selector maintenance, consistent with our findings of a dual input on CFI-1.

Different mechanisms are possible too, such as the “target reservoir” model for the terminal selector CHE-1 in ASE chemosensory neurons ^114^, or the “guarantor” model for ALR-1/ARX and HOX TFs in mechanosensory neurons ^107,112^. Alternatively, CFI-1, but not EGL-5, may exert a pioneering function that “opens up” closed chromatin surrounding “turned off” terminal identity genes, as its mammalian orthologs (ARID3) are known to affect chromatin structure ^115^ ^116^.

### Implications for systems neuroscience

Using the available *C. elegans* connectome^117^, we can make three specific predictions on information flow within the touch circuit. First, both CFI-1 and EGL-5 control the cholinergic identity of PVC cells (by activating *unc-17/VAChT*) and glutamatergic identity of LUA cells (by activating *eat-4/VGLUT*). Hence, loss of either TF affects the ability of PVC and LUA interneurons to respectively provide cholinergic and glutamatergic output to their post-synaptic targets (**Fig. 2C**). Second, loss of either TF also affects the transcription of multiple gap junction proteins (*inx-1, inx-11, inx-19, unc-7, unc-9*) - the building blocks of electrical synapses, which are prevalent in escape response circuits across species ^118,119^. Hence, the escape response defects in *cfi-1* or *egl-5* LOF mutants may arise, at least in part, due to impaired electrical transmission, consistent with a model where PVC electrical synapses facilitate escape responses to posterior touch stimuli (**Fig. 2C**)^12^. Third, both CFI-1 and EGL-5 control the expression of five glutamate (Glu) receptor-encoding genes (*glr-1, glr-2, glr-5, nmr-1, nmr-2*) in PVC interneurons. Hence, loss of either TF impairs the ability of PVC cells to receive input from glutamatergic mechanosensory (PVD, PLM) neurons (**Fig. 2C**).

Akin to reversible chemo- and opto-genetic approaches that manipulate neuronal activity by introducing exogenous genes (e.g., opsins, ion channels) ^120^, our study offers a blueprint to enable reversible, digital-like (ON, OFF) manipulations of adult neuron identity, circuit function, and animal behavior by tuning the expression levels of an endogenous TF-encoding gene.

### Limitations of the study

Because CFI-1 and EGL-5 control the expression of distinct sets of terminal identity genes in two different interneuron types, we hypothesize that additional TFs collaborate with CFI-1 and EGL-5 in either PVC or LUA cells. Although their identification is beyond the scope of the current study, experimental and computational work supports the notion that the LIM homeodomain protein CEH-14 and the Collier/Olf/Ebf (COE)-type TF UNC-3 control aspects of PVC identity ^47,121^. Future work is needed to test the hypothesis that CEH-14 and UNC-3 distinguish the cellular context of PVC from LUA, enabling CFI-1 and EGL-5 to activate distinct terminal identity genes in each interneuron type. Last, additional biochemical experiments are necessary to discern the precise molecular mechanism of how CFI-1 and EGL-5 bind DNA. Our motif proximity analysis showed that CFI-1 and EGL-5 motifs are separated by dozens of nucleotides, arguing against cooperative DNA binding as a heterodimer, a mechanism proposed for other terminal selector pairs^21,90^.

## ACKNOWLEDGEMENTS

We thank the Caenorhabditis Genetics Center (CGC), which is funded by NIH Office of Research Infrastructure Programs (P40 OD010440), for providing strains. We thank members of the Kratsios lab and Oliver Hobert for comments on the manuscript, as well as Jayson J. Smith for help with data presentation. This work was supported by a postdoc mobility fellowship (P500B-203088) from the Swiss National Science Foundation to F.M., a National Science Foundation predoctoral fellowship (Grant No. 2140001) to H.D., a Jeff Metcalf fellowship to M.M., and two NIH grants (R01 NS118078, R01NS116365) to P.K.

## AUTHOR CONTRIBUTIONS

**F**. M., Conceptualization, Data curation, Investigation, Visualization, Methodology, Writing— original draft, review and editing; Y.C., H. D., M. M., Formal analysis, Validation, Investigation, Writing—review and editing; P. K., Conceptualization, Supervision, Investigation, Funding acquisition, Project administration, Writing— original draft, review and editing.

## DECLARATION OF INTERESTS

The authors declare no competing interests.

## MATERIALS AND METHODS

### C. *elegans* strains and growth conditions

Worms were grown at 20 °C on nematode growth media (NGM) plates seeded with bacteria (OP50, Escherichia coli) as food source^122^. Single-mutant strains used in this study: OS122 *(cfi-1[ky651],* OH12344 *(cfi-1[ot786] I)*, MT1975 (*egl-5[n945] III*). All other strains used in this study are listed in **File S6**.

### Behavioral assays

L4 larvae were picked onto NGM plates one day before the experiments. Young adult worms were used for testing, unless otherwise indicated. All experimental replicates were performed on at least three independent days. Animal responses were scored as forward (green), backward (yellow), or no response (red).

#### Gentle touch

Gentle touch stimuli were delivered using an eyelash, as previously described ^123^. The stimulus was applied just behind the pharynx (anterior gentle touch) or near the tail region (posterior gentle touch) of non-moving animals.

#### Harsh touch

Harsh touch stimuli were delivered using a platinum wire pick, as previously described ^123^. The stimulus was applied from above by pressing the edge of the pick down just behind the pharynx (anterior harsh touch) or near the tail region (posterior harsh touch) of non-moving animals. To avoid any confounding effects of physical damage, each animal was tested only once.

### Generation of *C. elegans* transgenic animals

DNA constructs were generated using PCR fusion ^124^ and Gibson assembly. The constructs were then microinjected into the gonads of worms to generate transgenic lines, following a standard protocol ^125^. To identify transgenic animals, we used a [rol-6] co-injection marker.

### Cell-specific knockdown of *cfi-1*

We used the RNAi method based on the expression of sense and antisense RNA of a gene of interest, driven by cell-specific promoters ^76^. For LUA-specific RNAi, the *eat-4* and *plx-2* were used. For PVC-specific RNAi, the *nmr-1* promoter was used. Each promoter was fused to a 1 kb sense or antisense genomic sequence of *cfi-1*, encompassing exons 3 to 6. See **File S7** for primer sequences. The resulting PCR fusion DNA fragments, containing the sense or antisense *cfi-1* sequence, were co-injected together into young adult hermaphrodites carrying fluorescent reporters at 20 ng/µL each. A co-injection marker was used at 50 ng/µL. At least two independent transgenic lines were tested.

### Putative CFI-1 target genes in LUA and PVC interneurons

The top 1,000 highly expressed genes in LUA and PVC (measured in transcripts per million, TPM) at L4 were extracted from available scRNA-Seq data (CenGEN)^78^. This dataset was then computationally compared to a dataset of CFI-1 Chromatin Immunoprecipitation-sequencing (ChIP-seq) targets at L3 using the semi_join function in R (Dplyr package, version 1.0.7). This comparison generated a new data frame containing genes from the scRNA-seq dataset that are also putatively bound by CFI-1. Similarly, the set_diff function (Dplyr 1.0.7) was used to generate a data frame containing genes that are expressed in LUA or PVC based on scRNA-seq but are not found in the CFI-1 ChIP-seq dataset. Gene ontology analysis (Panther 19.0)^80^ was then performed on both data frames to functionally classify the genes based on protein class ontology.

### Transcription factor binding site analysis

The EGL-5 DNA binding site (motif) was obtained from EGL-5 ChIP-seq data from ENCODE (https://www.encodeproject.org/) (identifier: ENCFF440IOR) ^126^ ^127^, using Homer *de novo* motif analysis findMotifsGenome.pl, with parameters of peak size given and genome ce11^128^. The CFI-1 motif was obtained from CisBP, motif ID: M01749_3.00 ^129^. We have used these motifs to predict phylogenetically conserved CFI-1 and EGL-5 binding sites on the *C. elegans* genome using TargetOrtho2^79^, which can identify motifs in the upstream regions and introns of orthologous genes of up to eight nematode species.

### Inducible and reversible protein degradation with the AID system

AID-tagged proteins are conditionally degraded upon exposure to auxin in the presence of TIR1 ^52^. Animals carrying auxin-inducible alleles of *cfi-1 (kas16[mNG::AID::cfi-1])* or *egl-5 (syb2361 [egl-5::mNG::AID])* were crossed with *ieSi57* animals, which express TIR1 ubiquitously. Natural auxin, indole-3-acetic acid (IAA; Catalog number CAS 87-51-4, Alfa Aesar), was dissolved in ethanol (EtOH) to prepare a 400 mM stock solution. The synthetic auxin analog, K-NAA (Catalog number N610, PhytoTech Labs), was dissolved in distilled water (dH_2_O) to prepare a 400 mM solution. NGM plates containing 4 mM IAA or K-NAA (treatment) and EtOH or dH_2_O (control) were prepared, seeded with OP50 bacteria, and allowed to dry for 2-3 days ^53^.

#### Protein degradation

worms of the experimental strains were transferred onto auxin-treatment plates or control plates and kept at 20 °C for 1-3 days. All experimental plates were shielded from light.

#### Protein recovery

worms treated with 4mM auxin (IAA or K-NAA) and EtOH or dH_2_O were transferred onto standard NGM plates, incubated at 20 °C and allowed to recover for 1-5 days. After protein depletion or recovery, the worms were either imaged by fluorescence microscopy to monitor protein expression or subjected to behavioral assays to assess response to touch. All experimental replicates were obtained over at least three independent days.

### Microscopy

Worms were anesthetized with 100 mM of sodium azide (NaN_3_) and mounted on a 4% agarose pad on glass slides. Images were captured using an automated fluorescence microscope (Zeiss, Axio Imager.Z2) or a confocal microscope (Zeiss, LSM 900). Z-stack images (minimum thickness ∼1 μm) were acquired using the Zeiss Axiocam503 mono or LSM 900 systems. ZEN software, version 2.3.69.1000, Blue edition). Representative images are shown following maximum intensity projection of 10–20 μm Z-stacks. Image reconstruction was performed using Image J software^130^.

### Neurons identification

Neurons were identified based on a combination of the following criteria: (i) co-localization with fluorescent markers exhibiting known expression patterns, (ii) invariant cell body position and relative position to other neurons in the tail ganglia and (iii) use of the NeuroPAL strain ^71^.

### Fluorescence intensity quantification

To quantify fluorescence intensity (FI) in individual LUA and PVC interneurons, Z-stack images of the tail region were acquired with 0.5-1 μm intervals between stacks. At least 20 worms were analyzed per experimental condition or genotype. The same imaging parameters (e.g., exposure time, temperature) were applied across all samples. Image stacks were then processed (applied sliding paraboloid background subtraction method), and FI was quantified using FIJI ^130^. All neurons expressing the reporter were counted, and the FI in LUA and/or PVC cell bodies was quantified in arbitrary units (a.u).

### Statistical analysis and reproducibility

#### Behavioral graphs

Results are presented as percentage ^131^ and standard error of the mean (SEM; error bars). Significant differences between genotype or conditions are indicated as: *, p < 0.0001 versus WT; #, p < 0.0001 between mutants, as determined by Bonferroni post-hoc tests. In auxin experiments, *, p < 0.0001 versus control, #, p < 0.0001 versus auxin treatment.

#### Auxin experiments

For protein depletion, box and whisker plots were used to display all data points, with the whiskers extending to the minimum and maximum values, and the horizontal line within the box representing the median. Each dot corresponds to an individual neuron. This method also displays each individual value as a point superimposed on the graph. Statistical analyses were performed using an unpaired t-test (two-tailed) with Welch’s correction, and p-values were annotated. Differences with p < 0.01 were considered significant. For protein recovery, graphs show values expressed as mean ± standard deviation. Statistical analyses were performed using an unpaired t-test (two-tailed) with Welch’s correction and p-values were annotated as follows: *p < 0.05, **p < 0.01, ***p < 0.001.

#### Number of neurons graphs

Graphs show individual values expressed as mean ± standard error mean (SEM), with each dot representing an individual animal. Statistical analyses were performed using an unpaired t-test (two-tailed). Differences with p < 0.01 were considered significant. Asterisks in Fig. indicate statistical significance.

#### FI quantification graphs

For data quantification, box and whisker plots were used to display all data points, with the whiskers extending to the minimum and maximum values, and the horizontal line within the box representing the median. Each dot represents an individual neuron. This method also superimposes each individual value as a point on the graph. Statistical analyses were performed using an unpaired t-test (two-tailed) with Welch’s correction, and p-values were annotated. Differences with p < 0.01 were considered significant. Data visualization of data and p-value calculation were performed via GraphPad Prism Version 10.2.1.

## LEGENDS FOR SUPPLEMENTARY FIGURES, TABLES, AND FILES

**Figure S1:**
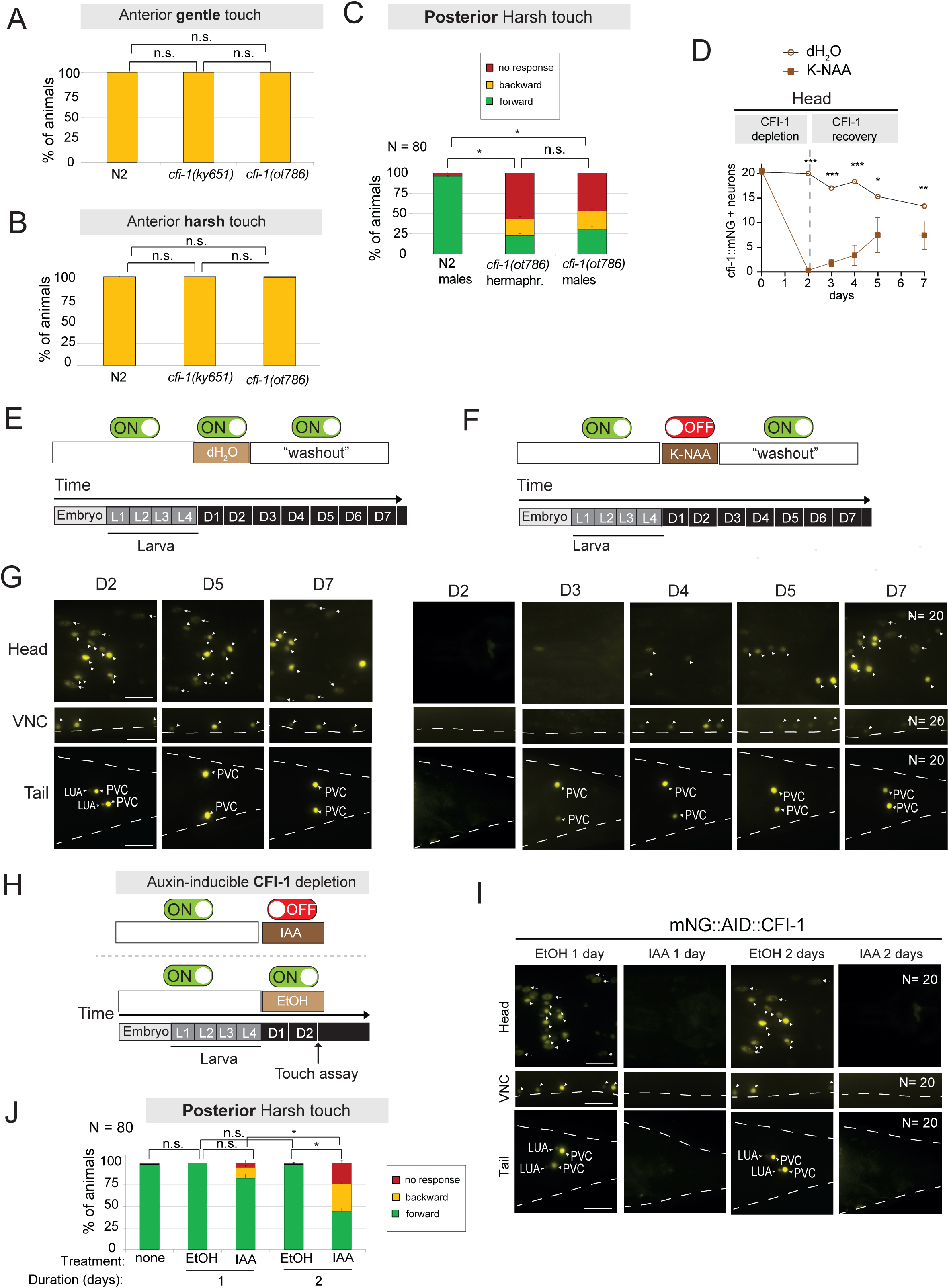
CFI-1 is required to maintain posterior harsh touch responses. **A**. Anterior gentle touch assay in *C. elegans* animals carrying *cfi-1* mutations. N = 80 animals. **B**. Anterior harsh touch assay in *C. elegans* animals carrying *cfi-1* mutations. N = 80 animals. **C**. Posterior harsh touch assay in *cfi-1(ot786)* mutant animals. Comparison between hermaphrodites and males. N = 80 animals. **D**. Quantification of head neurons expressing the endogenous CFI-1 reporter after water (control), auxin and “washout” recovery treatments on *kas16[mNG::AID::cfi-1]; ieSi57 [eft-3p::TIR1::mRuby::unc-54 3’UTR + Cbr-unc-119(+)]* animals. N = 5 animals. **E**. Diagrams illustrating the timeline of the auxin (K-NAA) and “washout” treatments used for the depletion of CFI-1. **F**. Diagrams illustrating the timeline of the auxin (K-NAA) and “washout” treatments. **G**. Recovery of mNG::AID::CFI-1 expression upon auxin treatment followed by washout. Representative images showing mNG::AID::CFI-1 expression in the head, ventral nerve cord (VNC) and tail of *C. elegans* animals. Arrowheads point to neurons, while the arrows point to head muscle cells. N = 20 animals. **H**. Diagram illustrating the timeline of the EtOH (control) and auxin (IAA) treatments used for the depletion of CFI-1. **I**. Representative images showing the loss of mNG::AID::CFI-1 expression in the head, VNC and tail of *C. elegans* animals. Arrowheads point to neurons, while the arrows point to head muscle cells. N = 20 animals. **J**. Posterior harsh touch assay in *C. elegans* animals before and after EtOH (control) and auxin (IAA) treatments for 1 and 2 days. N = 80 animals. For behavior in panels **A**, **B**, **C** and **J**: *, p < 0.0001; n.s., not significant, as determined by Bonferroni post-hoc tests. For quantifications in panel **D**: *, p = 0.002; **, p = 0.00003; ***, p < 0.000003 versus auxin treatment, as determined by Bonferroni post-hoc test.

**Figure S2:**
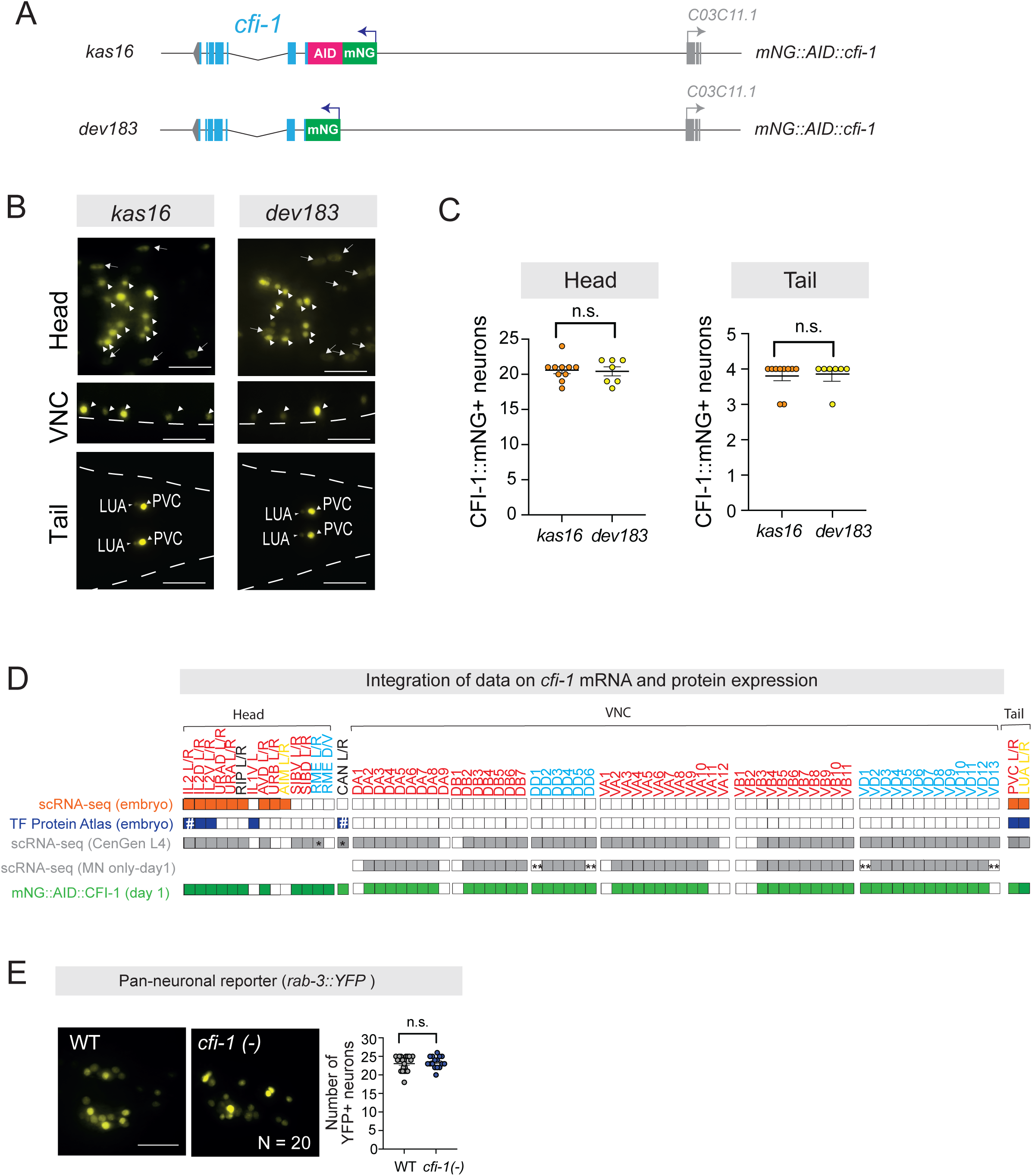
CFI-1 expression analysis using endogenous reporters and available RNA-Seq datasets. **A**. Diagram showing the two *cfi-1* endogenous reporter alleles. The *kas16* allele is described in Figure 1; *dev183* allele: *cfi-1* endogenous reporter with an in-frame fluorescent protein mNeonGreen (mNG) insertion. **B**. Representative images showing CFI-1 expression in the head, ventral nerve cord, and tail of *kas16* and *dev183* adult animals. Arrowheads point to neurons, while the arrows point to head muscle cells. N = 10 animals. **C**. Graphs showing the number of head and tail neurons expressing either CFI-1 endogenous reporters. **D**. Comparison of endogenous *cfi-1* expression from embryo to adulthood in different studies. *, represents low value transcript per million; #, means that expression was detected only in IL2L and CANL; **, neurons were not isolated in that study. **E**. Expression analysis of a fluorescent reporter for a pan-neuronal marker *rab-3* in WT and *cfi-1(ot786)* mutant young adult animals. N = 20 animals. For panels **C** and **E**, scatter dot plots displaying number of neurons expressing respective fluorescent reporter, with the horizontal line representing the mean with standard error of the mean (SEM). Each dot represents the average number of reporter (+) neurons in one animal. An unpaired *t*-test (two-sided) with Welch’s correction was performed: n.s., not significant between *cfi-1* reporters or versus WT. Scale bars: 20 μm.

**Figure S3:**
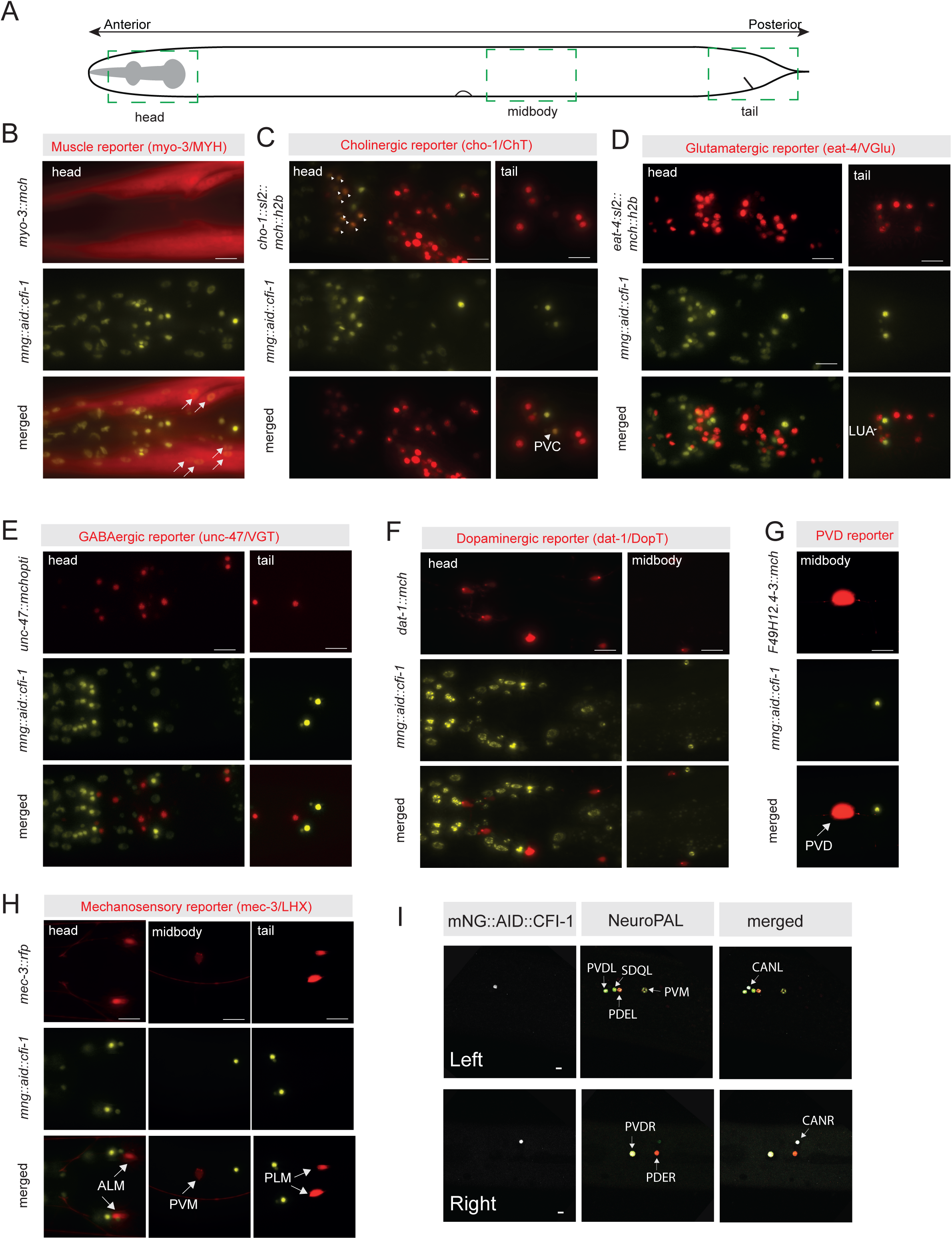
CFI-1 expression analysis using molecular markers. **A**. Schematic showing a lateral view of an entire worm, with dashed boxes highlighting the regions shown in the images below **B**. Representative images of the worm head region showing CFI-1 expression in combination with a muscle reporter (*myo-3*). CFI-1 is expressed in head muscle cells. Arrows point to head muscle cells. N = 10 animals. **C**. Representative images of the worm head and tail regions showing CFI-1 expression in combination with a cholinergic neuron reporter (*cho-1*). CFI-1 is expressed in approximately 13 cholinergic neurons in the head and 2 cholinergic neurons in the tail. Arrowhead points to PVC interneuron. N = 10 animals. **D**. Representative images of the worm head and tail regions showing CFI-1 expression in combination with a glutamatergic neuron reporter (*eat-4*). CFI-1 is expressed in 2 glutamatergic neurons in the tail. Arrowhead points to LUA interneuron. N = 10 animals. **E**. Representative images of the worm head and tail regions showing CFI-1 expression in combination with GABAergic neuron reporter (*unc-47*). CFI-1 in not expressed in any GABAergic neurons. N = 10 animals. **F**. Representative images of the worm head and posterior midbody regions showing CFI-1 expression in combination with a dopaminergic reporter (*dat-1*). CFI-1 does is not expressed in any dopaminergic neuron. N = 10 animals. **G**. Representative images of the worm posterior midbody region showing CFI-1 expression in combination with a specific PVD-specific sensory neuron reporter. CFI-1 is not expressed in PVD sensory neurons. Arrow points to PVD neuron. N = 10 animals. **H**. Representative images of the worm head, midbody and tail regions showing CFI-1 expression in combination with a mechanosensory reporter (*mec-3*). CFI-1 is not expressed in any mechanosensory neurons. Arrows point to mechanosensory neurons (ALM/PLM/PVM). N = 10 animals. **I**. Representative images of the posterior midbody showing CFI-1 expression in the CAN neurons, in conjunction with the NeuroPAL strain. Scale bars: 20 μm.

**Figure S4:**
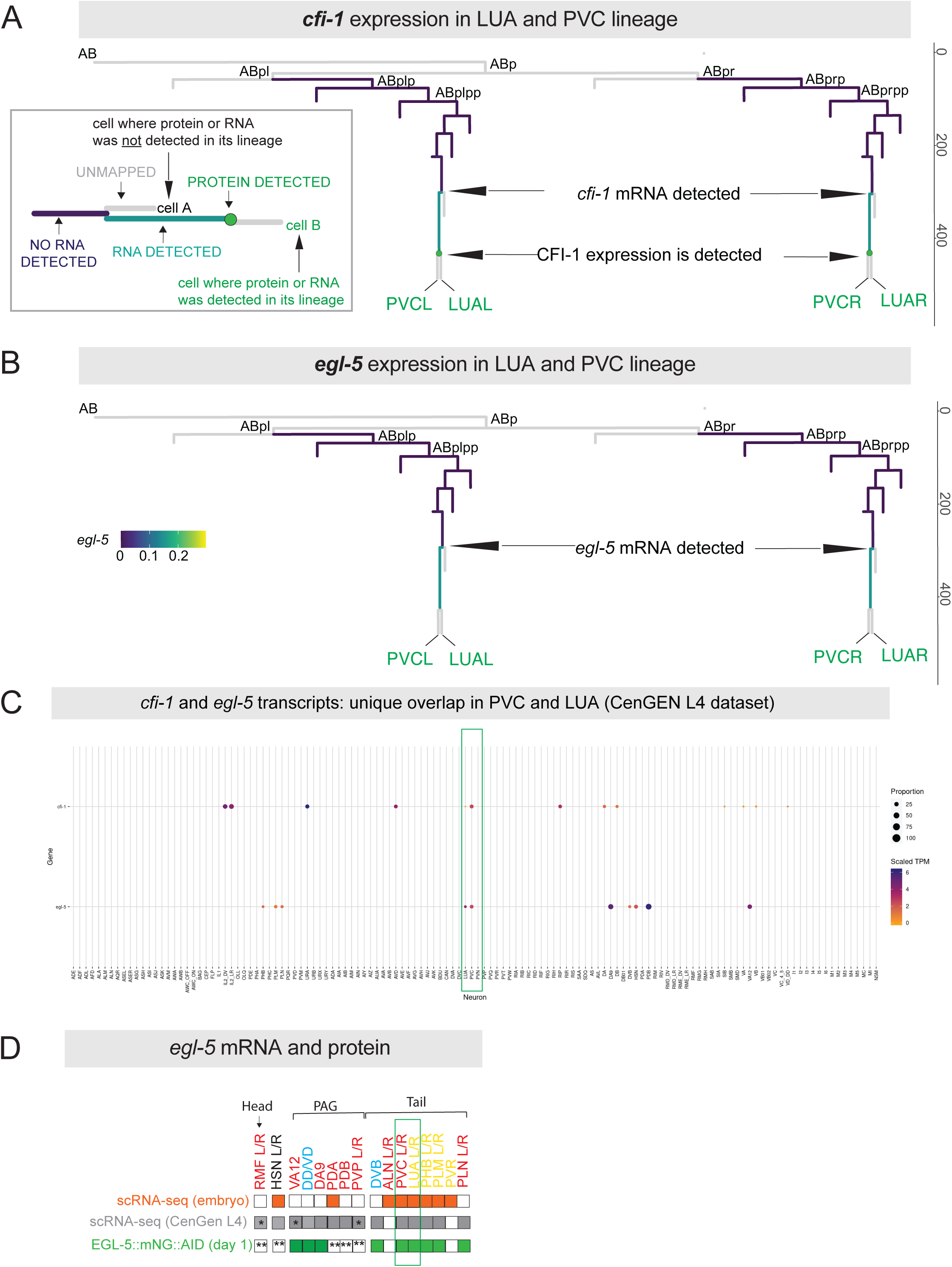
*cfi-1* and *egl-5* mRNA transcripts are detected in the mother cell of PVC and LUA sister interneurons, but not earlier in the lineage. **A-B.** Cell lineage trees of *C. elegans* from the embryonic stage to the bean stage showing the expression of *cfi-1* (**A**) and *egl-5* (**B**) in LUA and PVC lineage. This tree combines scRNA data from Parker et al (2019) and protein expression data from the Ma et al (2021) protein atlas. (Bottom left) (**A**) Diagram explaining the lineage tree. **C.** Heatmap of *cfi-1* and *egl-5* expression showing relative expression levels and proportion of neurons expressing these genes. The color code represents, scaled expression of *cfi-1* and *egl-5* (from CeNGEN), with the green box highlighting LUA and PVC interneurons. **D.** Comparison of endogenous *egl-5* expression from embryo to adulthood in different studies. *, represents low value transcript per million; **, not determined (n.d.).

**Figure S5:**
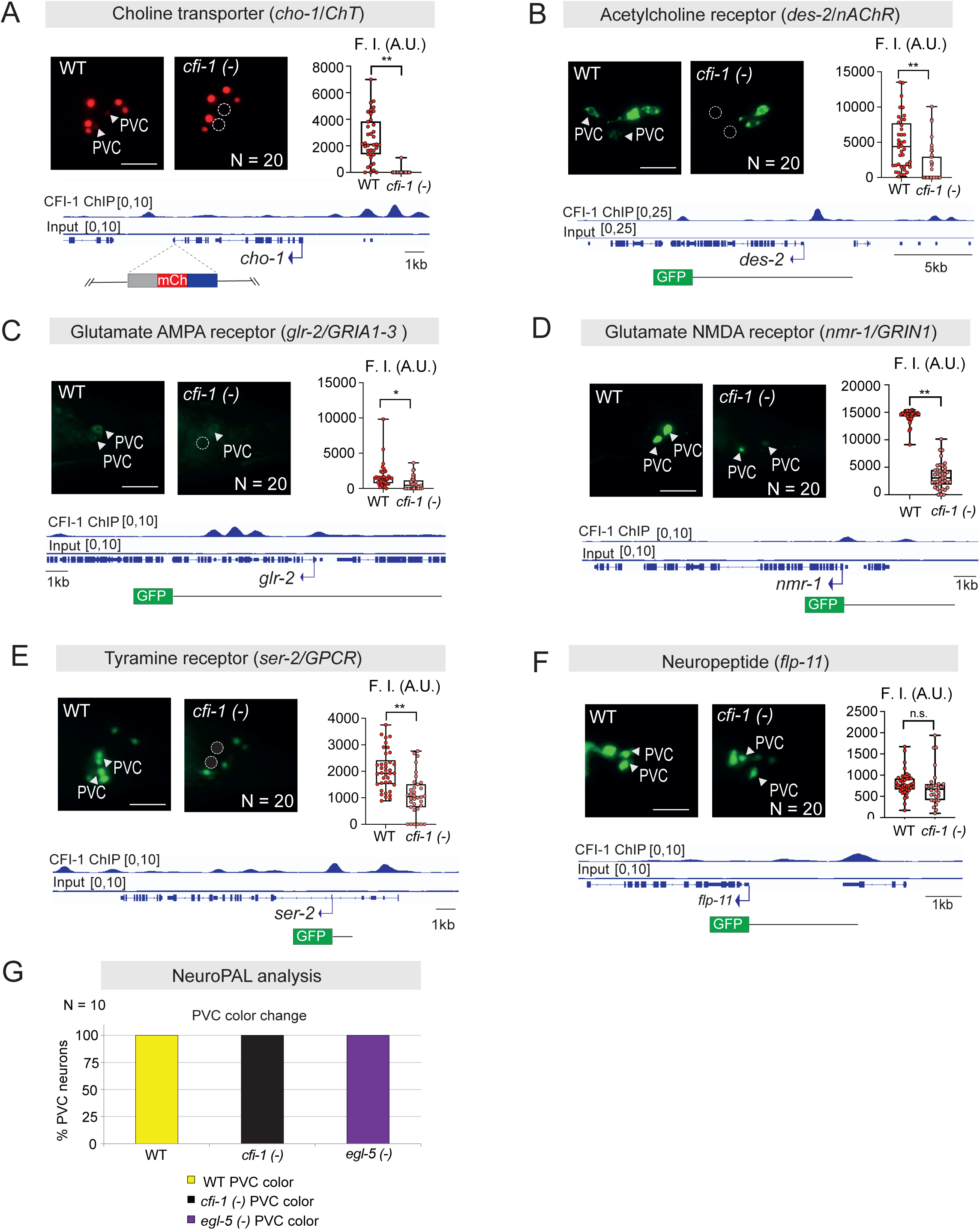
CFI-1 activates multiple terminal identity genes in PVC interneurons. **A-F**. Expression analysis of six fluorescent reporters of PVC terminal identity (*cho-1, des-2, glr-2, nmr-1, ser-2* and *flp-11*) in WT and *cfi-1(ot786)* mutant young adult animals. N = 20 animals. **G.** Graph showing the percentage of PVC interneurons that change color in either the *cfi-1* or *egl-5* mutant backgrounds, compared to WT. For panels **A-F**, (Top left) Representative images showing reporters expression. White arrowheads indicate expression in PVC interneurons, while the white dashed circle marks the absence of expression in PVC interneurons. (Bottom) IGV snapshots showing CFI-1 binding at loci of genes, along with a diagram of the reporter allele used. (Top right) Box and whisker plots displaying quantitative fluorescence intensity (F.I) measurements in arbitrary units (A.U.), with the whiskers extending to the minimum and maximum values, and the horizontal line within the box representing the median. Each dot represents an individual neuron. An unpaired *t*-test (two-sided) with Welch’s correction was performed: n.s., non-significant; *, p = 0.0028; **, p < 0.0001 versus WT. Scale bars: 20 μm.

**Figure S6:**
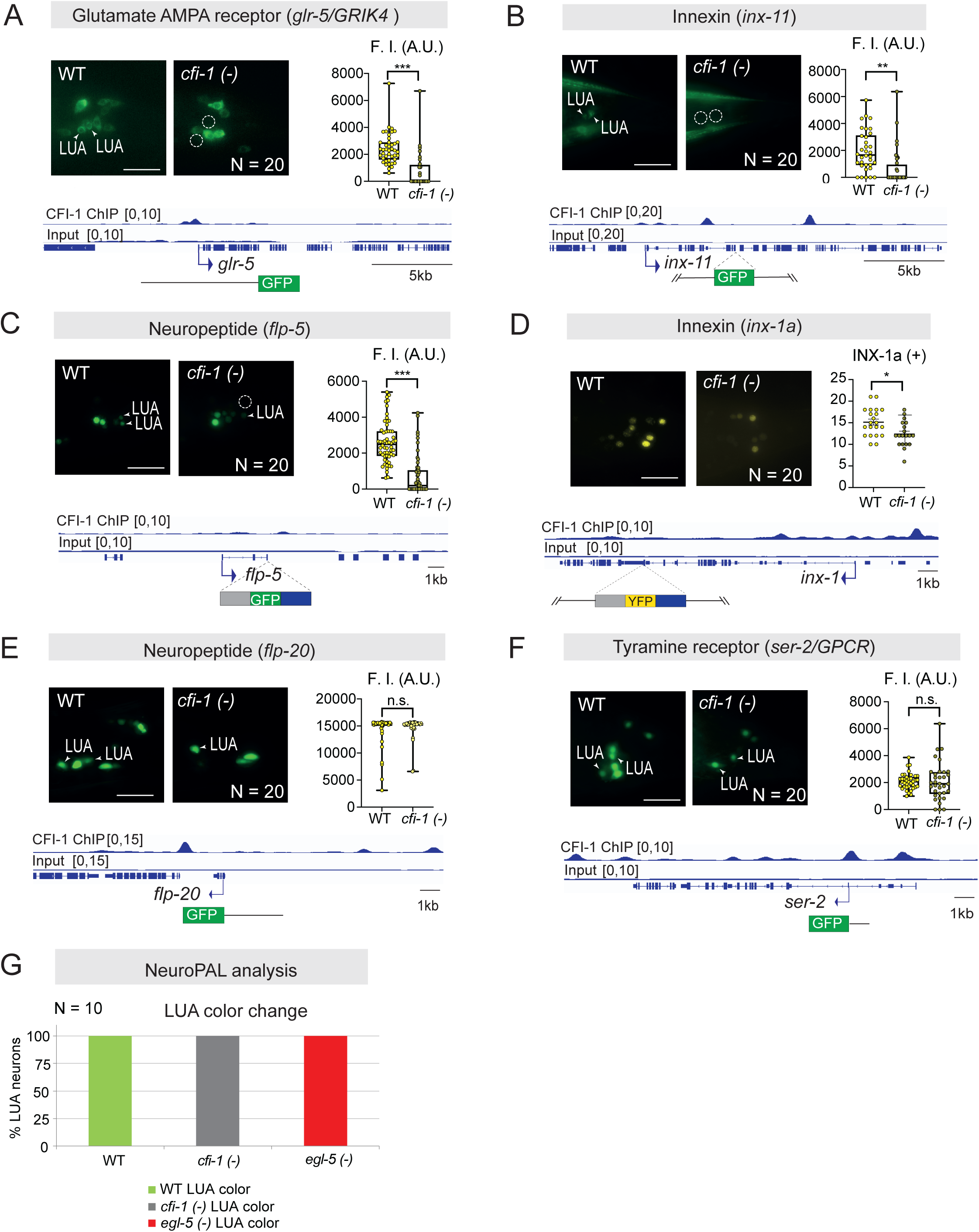
CFI-1 activates multiple terminal identity genes in LUA interneurons. **A-F**. Expression analysis of six fluorescent reporters of LUA terminal identity (*glr-5, inx-11, flp-5, inx-1, flp-20,* and *ser-2*) in WT and *cfi-1(ot786)* mutant young adult animals. N = 20 animals. **G.** Graph showing the percentage of LUA interneurons that change color in either *cfi-1* or *egl-5* mutant backgrounds, compared to WT. For panels **A-F**, (Top left) Representative images showing reporters expression. White arrowheads indicate expression in LUA interneurons, while the white dashed circle marks the absence of expression in LUA interneurons. (Bottom) IGV snapshots showing CFI-1 binding at loci of genes, along with a diagram of the reporter allele used. (Top right) Box and whisker plots displaying quantitative fluorescence intensity (F.I) measurements in arbitrary units (A.U.), with the whiskers extending to the minimum and maximum values, and the horizontal line within the box representing the median. Each dot represents an individual neuron. An unpaired *t*-test (two-sided) with Welch’s correction was performed: n.s., non-significant; *, p = 0.0072; **, p = 0.0001, ***, p < 0.0001 versus WT. In panel **D**, *inx-1* gene is also expressed in PVC interneurons. Scale bars: 20 μm.

**Figure S7:**
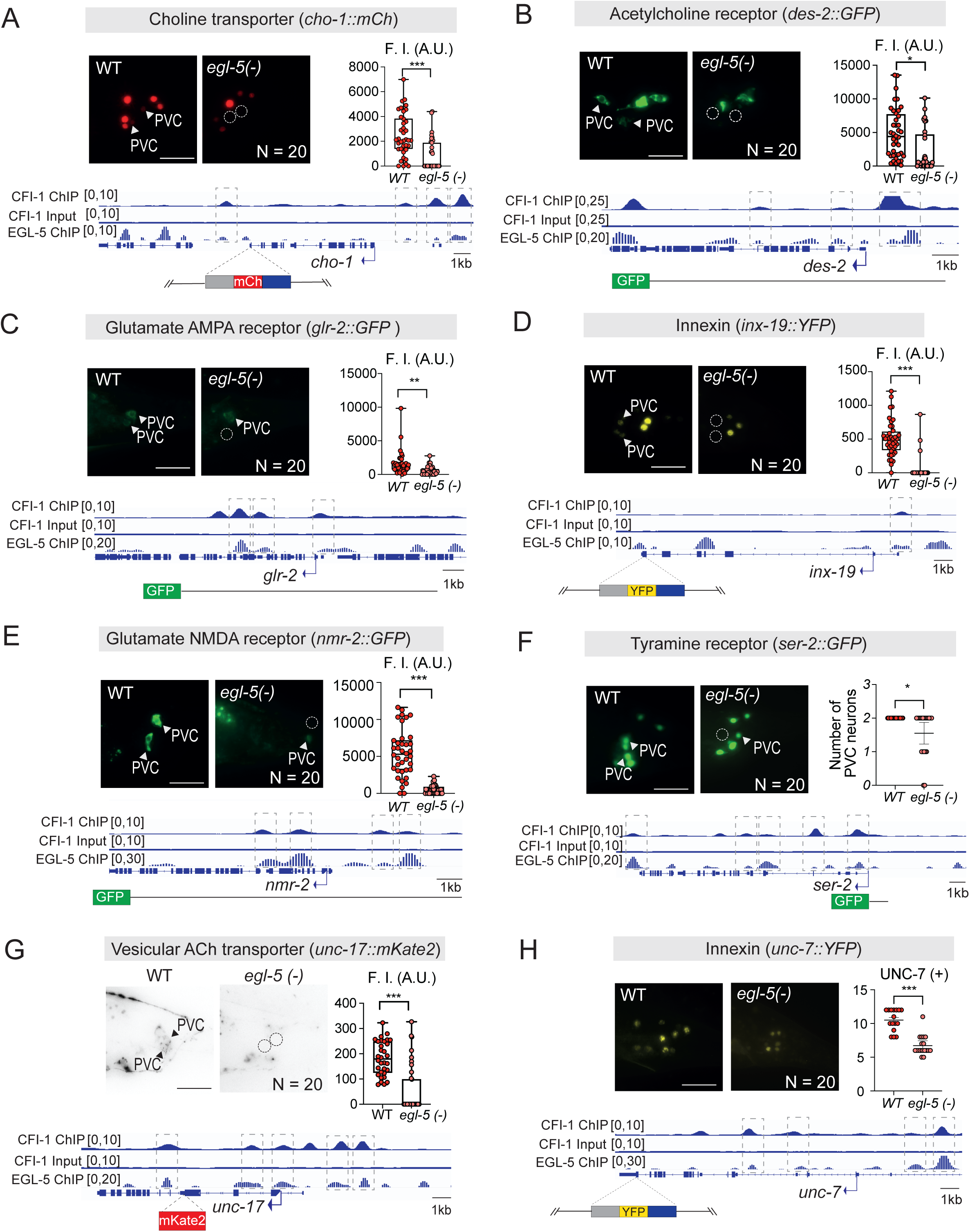
EGL-5 activates multiple terminal identity genes in PVC interneurons. **A-H**. Expression analysis of eight fluorescent reporters of PVC terminal identity (*cho-1, des-2, glr-2, inx-19, nmr-2, ser-2, unc-17* and *unc-7*) in WT and *egl-5(n945)* mutant young adult animals. N = 20 animals. (Top left) Representative images showing reporters expression. WT image shown for *egl-5* analysis is the same as in *cfi-1* mutant analysis. White arrowheads indicate expression in PVC interneurons, while the white dashed circle marks the absence of expression in PVC interneurons. (Bottom) IGV snapshots showing CFI-1 and EGL-5 binding at *loci* of genes and a diagram of the reporter allele used. Dashed boxes highlight the CFI-1 and EGL-5 ChIP peaks that are in the same region. EGL-5 input track is not shown because original authors normalized to the EGL-5 ChIP signal. (Top right) Box and whisker plots displaying quantitative fluorescence intensity (F.I) measurements in arbitrary units (A.U.), with the whiskers extending to the minimum and maximum values, and the horizontal line within the box representing the median. Each dot represents an individual neuron. (F) Scatter dot plot displaying number of PVC neurons expressing *ser-2* reporter, with the horizontal line representing the mean with standard error of the mean (SEM). Each dot represents the average number of reporter (+) PVC neurons in one animal. An unpaired *t*-test (two-sided) with Welch’s correction was performed: (**B**) *, p = 0.002; (**C**) **, p = 0.0005;; (**F**) *, p = 0.009 versus WT; (**A,D, E,G,H**) ***, p < 0.0001 versus WT. Scale bars: 20 μm.

**Figure S8:**
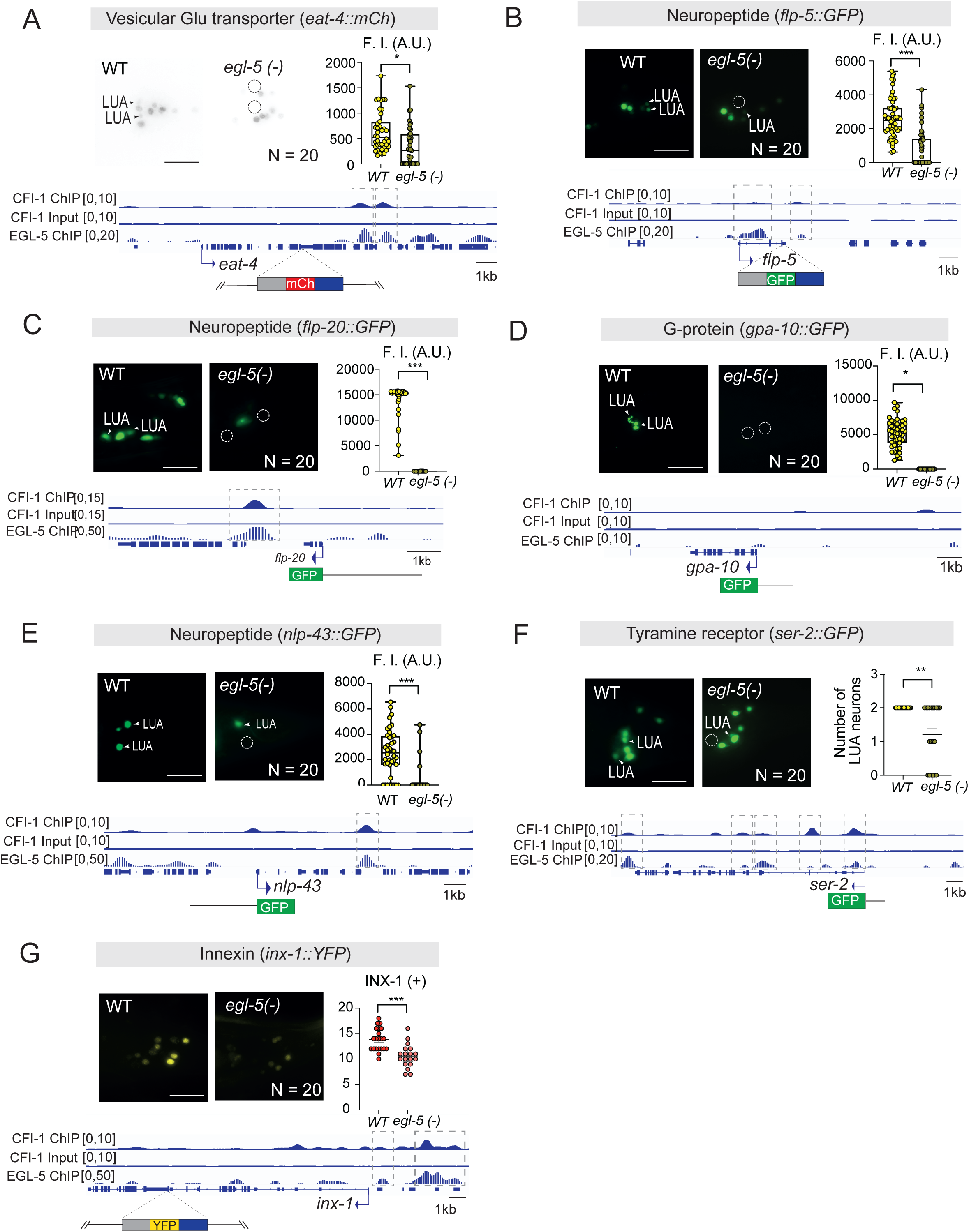
EGL-5 acts as terminal selector in LUA interneurons. **A-G**. Expression analysis of seven fluorescent reporters of LUA terminal identity (*eat-4, flp-5, flp-20, gpa-10, nlp-43, ser-2,* and *inx-1*) in WT and *egl-5(n945)* mutant young adult animals. N = 20 animals. (Top left) Representative images showing reporters expression. WT image shown for *egl-5* analysis is the same as in *cfi-1* mutant analysis. White arrowheads indicate expression in LUA interneurons, while the white dashed circle marks the absence of expression in LUA interneurons. (Bottom) IGV snapshots showing CFI-1 and EGL-5 binding at *loci* of genes and a diagram of the reporter allele used. Dashed boxes highlight the CFI-1 and EGL-5 ChIP peaks that are in the same region. EGL-5 input track is not shown because original authors normalized to the EGL-5 ChIP signal. (Top right) Box and whisker plots displaying quantitative fluorescence intensity (F.I) measurements in arbitrary units (A.U.), with the whiskers extending to the minimum and maximum values, and the horizontal line within the box representing the median. Each dot represents an individual neuron. (F) Scatter dot plot displaying number of LUA neurons expressing *ser-2* fluorescent reporter, with the horizontal line representing the mean with standard error of the mean (SEM). Each dot represents the average number of reporter (+) LUA neurons in one animal. An unpaired *t*-test (two-sided) with Welch’s correction was performed: (**A**) *, p = 0.001; (**D**) *, p = 0.002; (**F**) **, p = 0.0008 versus WT; (**B,C,E,G**) ***, p < 0.0001 versus WT. Scale bars: 20 μm.

**Figure S9:**
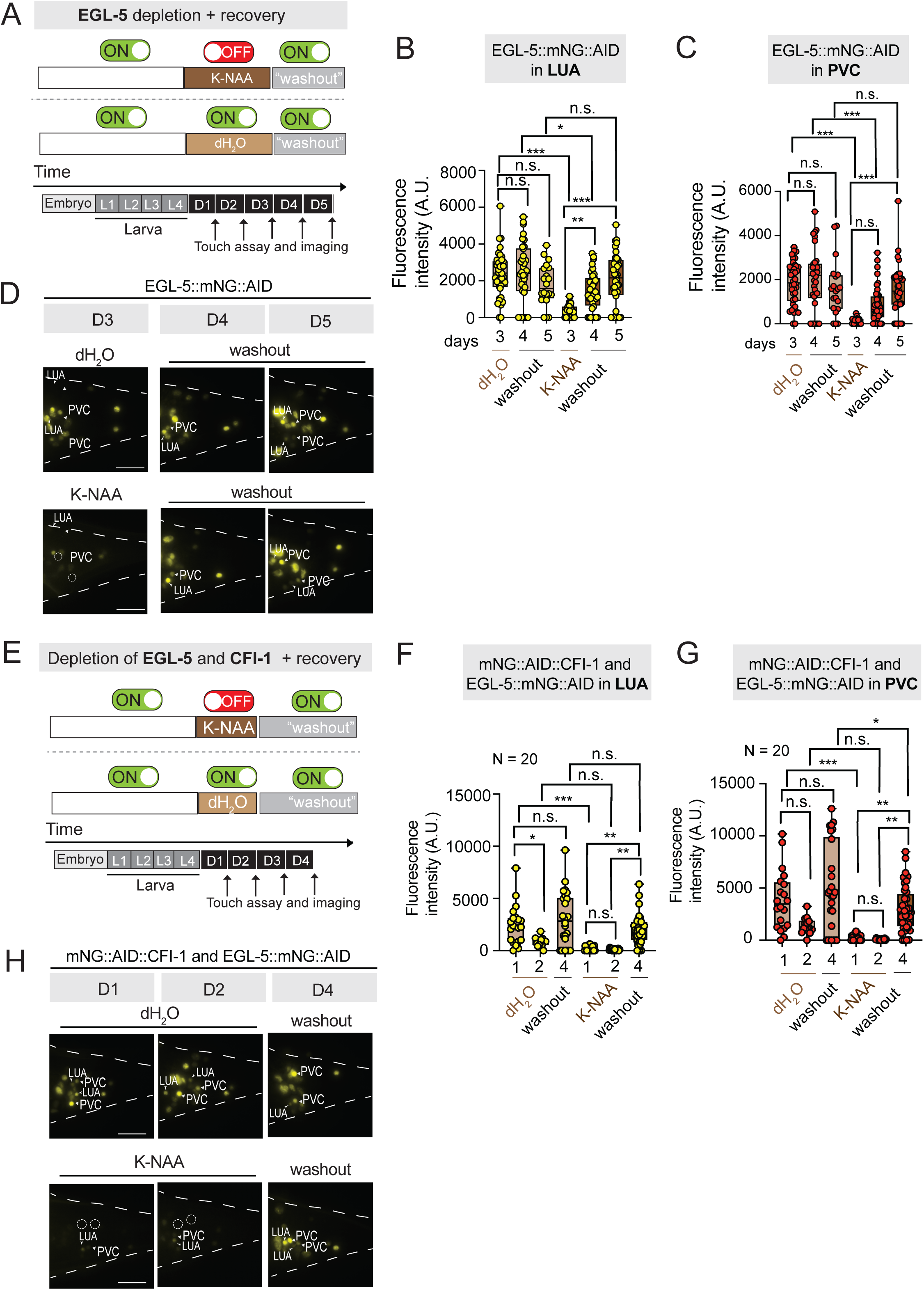
CFI-1 and EGL-5 expression can be restored after depletion in the adult. **A.** Diagrams illustrating the timeline of the auxin (K-NAA) and “washout” treatments used for the depletion of EGL-5. **B-C.** Quantification of LUA (B) and PVC (C) neurons expressing the endogenous EGL-5 reporter after water (control), auxin and “washout” recovery treatments on *syb2361 [egl-5::mNG::AID]; ieSi57 [eft-3p::TIR1::mRuby::unc-54 3’UTR + Cbr-unc-119(+)]* animals. N = 10 animals. **D.** Recovery of EGL-5::mNG::AID expression upon auxin treatment followed by washout. Representative images showing EGL-5::mNG::AID expression in tail of *C. elegans* animals. N = 10 animals. **E.** Diagrams illustrating the timeline of the auxin (K-NAA) and “washout” treatments used for the simultaneous depletion of CFI-1 and EGL-5. **F-G.** Quantification of LUA (B) and PVC (C) neurons expressing the endogenous CFI-1 and EGL-5 reporters after water (control), auxin and “washout” recovery treatments on *kas16[mNG::AID::cfi-1]*; *syb2361 [egl-5::mNG::AID]; ieSi57 [eft-3p::TIR1::mRuby::unc-54 3’UTR + Cbr-unc-119(+)]* animals. N = 10 animals. **H.** Recovery of mNG::AID::CFI-1 and EGL-5::mNG::AID expression upon auxin treatment followed by washout. Representative images showing mNG::AID::CFI-1 and EGL-5::mNG::AID expression in tail of *C. elegans* animals. N = 20 animals. For panels **B-C** and **F-G,** box and whisker plots are displaying quantitative fluorescence intensity (F.I) measurements in arbitrary units (A.U.), with the whiskers extending to the minimum and maximum values, and the horizontal line within the box representing the median. Each dot represents an individual neuron. An unpaired *t*-test (two-sided) with Welch’s correction was performed: *, p = 0.0038; **, p = 0.0010 (**B**); *, p = 0.0261; **, p = 0.0003 (**F**); *, p = 0.0259; **, p = 0.0004 (**G**); ***, p < 0.0001; n.s. not significant (**B-C, F-G**) as determined by Bonferroni post-hoc test. Scale bars: 20 μm.

**Figure S10:**
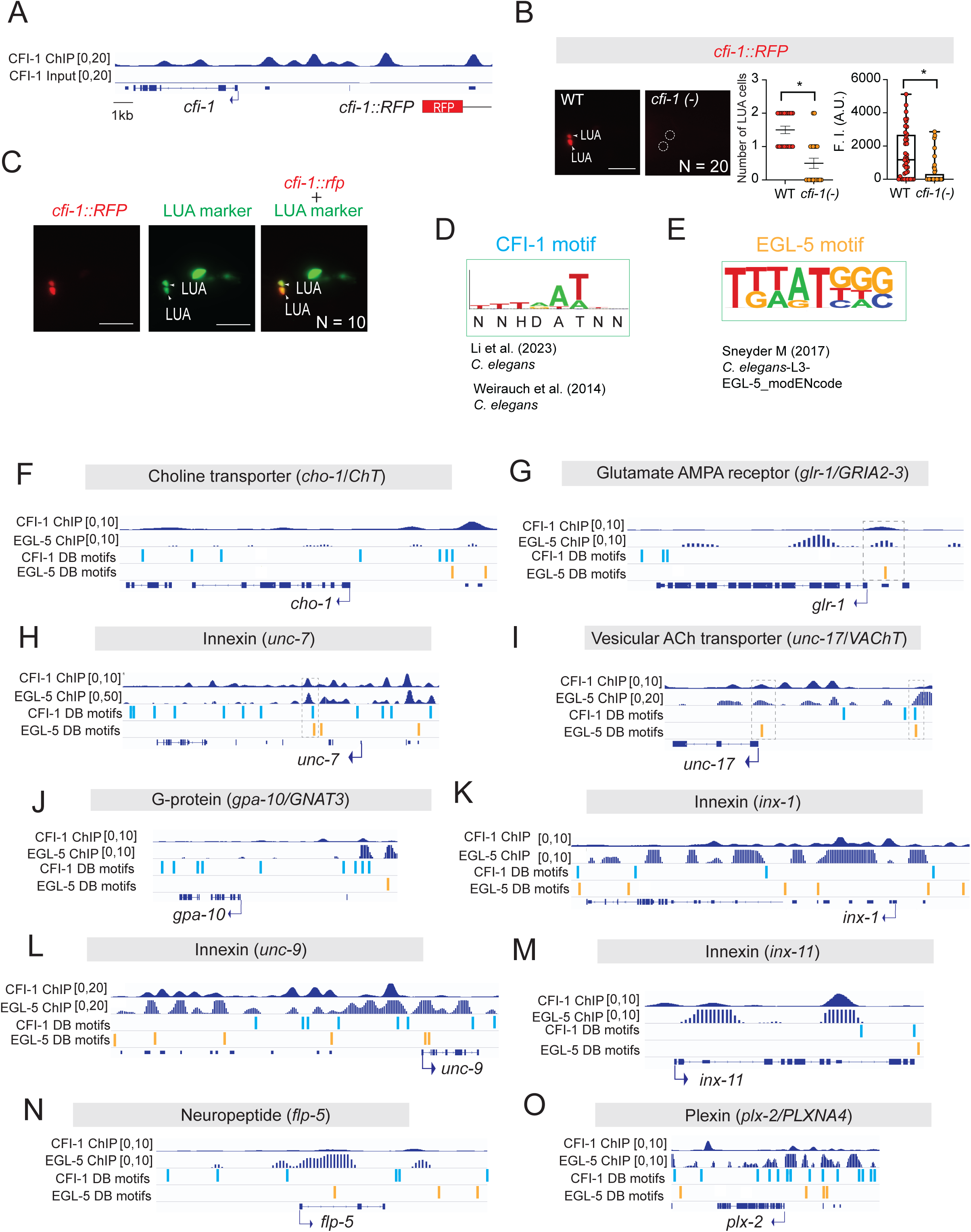
CFI-1 and EGL-5 DNA binding motifs are well conserved across species. **A**. IGV snapshot showing CFI-1 binding at its own locus, alongside a diagram of the reporter allele used. **B**. Expression analysis of *cfi-1*::RFP in WT and *cfi-1(ot786)* mutant young adult animals, N = 20 animals. **C**. Representative images of the worm tail region showing *cfi-1p*::RFP expression in combination with a LUA reporter (*eat-4::gfp*). White arrowheads indicate expression in LUA interneurons. N = 10 animals. **D**. DNA binding motif of CFI-1. **E**. de novo motif discovery analysis of EGL-5 binding peaks identifies an 8 bp-long EGL-5 binding motif. (**F-O**). IGV snapshots showing CFI-1 and EGL-5 binding at *loci* of genes together with their DB motifs found by TargetOrtho 2 analysis. Dashed boxes highlight the CFI-1 and/or EGL-5 DB motifs that are in the same region of the ChIP peaks.

**Table S1: A nervous system-wide map of *cfi-1/ARID3* expression in *C. elegans***

**Table S2. Summary of CFI-1 and EGL-5 target genes in PVC interneurons**

**Table S3. Summary of CFI-1 and EGL-5 target genes in LUA interneurons**

**File S1: CFI-1 ChIP-Seq analysis on top 1,000 genes expressed in PVC and LUA interneurons.**

**File S2: TargetOrtho2 analysis of CFI-1 and EGL-5 motifs.**

**File S3: GO analysis on top 1,000 genes expressed in PVC interneurons**

**File S4: GO analysis on top 1,000 genes expressed in LUA interneurons**

**File S5: Validation of PVC and LUA markers**

**File S6: List of strains used in this work**

**File S7: List of primers used in this work**

## Notes

### Competing Interest Statement

The authors have declared no competing interest.

